# Two antagonistic effect genes mediate separation of sexes in a fully dioecious plant

**DOI:** 10.1101/2020.03.15.993022

**Authors:** Liangjiao Xue, Huaitong Wu, Yingnan Chen, Xiaoping Li, Jing Hou, Jing Lu, Suyun Wei, Xiaogang Dai, Matthew S. Olson, Jianquan Liu, Mingxiu Wang, Deborah Charlesworth, Tongming Yin

**Affiliations:** The Key Laboratory of Tree Genetics and Biotechnology of Jiangsu Province and Education Department of China, Nanjing Forestry University, Nanjing, China, 200137; Department of Biological Sciences, Texas Tech University, Lubbock, TX 79409, USA; Key Laboratory of Bio-Resource and Eco-Environment of Ministry of Education, College of Life Sciences, Sichuan University, Chengdu 610065, China; Institute of Evolutionary Biology, University of Edinburgh, Charlotte Auerbach Road, Edinburgh EH9 3FL, UK

## Abstract

Plant sex determining systems and sex chromosomes are often evolutionarily young. Here, we present the early stage of sex chromosome in a fully dioecious plant, *P. deltoides*, by determining separate sequences of the physically small X- and Y-linked regions. Intriguingly, two Y genes are absent from the X counterpart. One gene represses female structures by producing siRNAs that block expression of a gene necessary for development of female structures, via RNA-directed DNA methylation and siRNA-guided mRNA cleavage. The other gene generates long non-coding RNA transcripts that, in males, soak up miRNAs that specifically inhibit androecium development. Transformation experiments in *Arabidopsis thaliana* show that the two genes affect gynoecium and androecium development independently and antagonistically. Sex determination in the poplar therefore has the properties proposed for the first steps in the evolution of dioecy in flowering plants, with two genes whose joint effects favor close linkage, as is observed in poplar.

---

Sex determination is an interesting aspect of plant reproduction, development and evolution (*1-3*). Most flowering plants produce so called “perfect” bisexual flowers (hermaphroditism). Only about 10% of angiosperms bear unisexual flowers, either with male and female flowers on the same plant (monoecism) or on separate individuals (dioecism) (*2*). Dioecism has evolved independently hundreds of times from hermaphroditic ancestors, in multiple plant lineages (*2*), and recent advances have allowed details of its genetic control to be understood in several dioecious plants (*e.g.* (*4-7*). A theoretical model for the origin of sex chromosomes involves a transition from functional hermaphroditism (including monoecy) to dioecism via mutations in two linked genes acting independently and antagonistically on female and male functions (*8, 9*). Recently, empirical studies in garden asparagus (*Asparagus officinalis* L.) (*7*) and kiwifruit (*Actindia rufa* × *A. chinensis*) (*10, 11*), have supported this hypothesis, by revealing two fully Y-linked genes that trigger the developmental pathway leading to males, rather than females (*6*).

A two-gene system is not inevitable, single-gene sex determination, involving a single gene that dominantly suppresses female function and promotes male function can be experimentally created in monoecious plants, including maize (*12*) and melon (*Cucumis melo*) (*13*), by fixing null mutations in unlinked but interacting genes, one of which acts in a sex-specific manner. Sex determination in the persimmon (*Diospyros lotus*, a tree) is a naturally evolved single-gene system, involving a non-coding RNA locus, *OGI*, suppressing femaleness (*6*). However, two mutations were necessary for its evolution, and the target gene *MeGI* is also inferred to have changed during the evolution of females (*14*).

Given the independent origins of dioecism in flowering plants, different genetic factors are likely to control sex determination in unrelated plant lineages (*7*). Unlike the “cryptic dioecy” in garden asparagus and kiwifruit, whose flowers bear apparently normal organs of the opposite sex, sex organ abortion in poplars occurs early, before the initiation of stamen or carpel primordia. Poplar chromosome XIX carries the sex determining locus (*15-21*), and a number of candidate genes have been proposed (*22-26*). In this study, we cloned the sex determination genes in *P. deltoides*, characterized their regulation mechanisms and functions using multiple approaches, and suggest a pathway by which dioecy could have evolved.

## Results

### Mapping the sex-determining locus and reconstructing X and Y haplotypes

Linkage analysis using simple sequence repeat (SSR) markers (Supplementary Table 1) located the sex-determination locus to the peritelomeric end of chromosome XIX (Supplementary Fig. 1), and its segregation confirmed male heterogamety (XY sex determination system). We sequenced and *de novo* assembled the genomes of a poplar female and one of her male offspring. The assembly for the female is 431 Mb, with contig N50 of 1.4 Mb, and 414 Mb with contig N50 of 2.8 Mb for the male. Our SSR markers located the sex determination locus to a 299 kb region between the end of the telomeric region and the N362 marker (Fig. 1A), while the centromere-proximal region recombines, and is pseudo-autosomal (PAR); we refer to the region terminal to N362 as the sex-linked region (SLR).

**Fig. 1.**
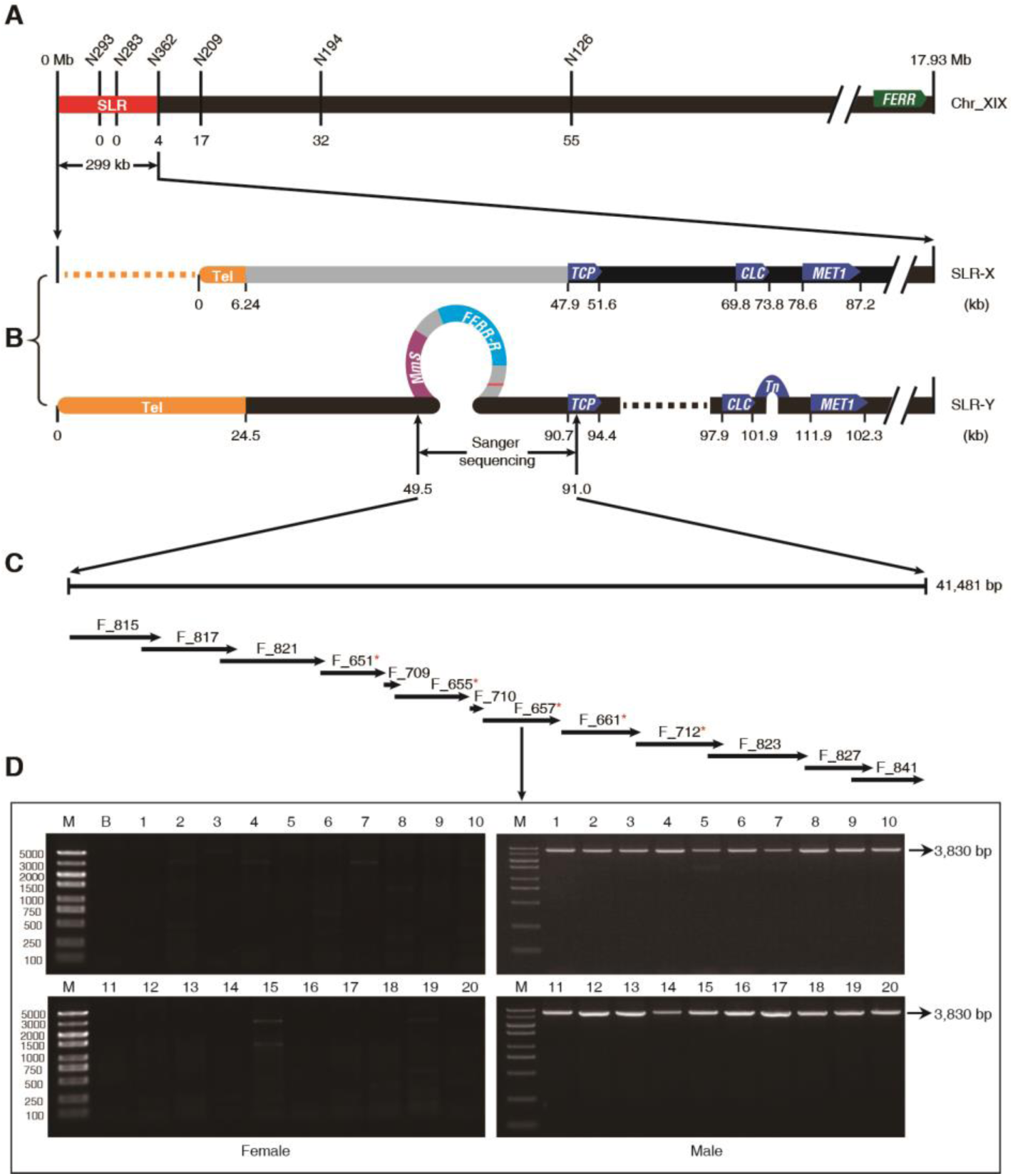
Reconstructed haplotypes in the SLR of *P. deltoides* chromosome XIX. (**A**) Genetic map and physical positions of the SLR (red bar) and the telomere. SSR makers are shown on the top and the numbers of observed recombination events are shown under the chromosome diagram, with zero in both the genetic and physical maps at the telomere end of the chromosome. The *FERR* gene is located at the right end of chromosome XIX, the end most distant from the telomere of this telocentric chromosome. (**B**) SLR-X and SLR-Y haplotypes reconstructed from genome sequences of our sequenced male (see main text). The yellow bars at the left indicate the X and Y telomeres, and physical distances (kb) from the telomere end are shown under each bar. The dashed lines represent deleted sequences, whereas the loops on SLR-Y represent the two Y-specific fragments (YHF) described in the text. The gray portion on SLR-X indicates a region described in the text, where divergence between SLR-X and SLR-Y is higher than elsewhere in the SLR. (**C**) The Y-specific region rebuilt from 13 PCR amplified fragments (fragment names are shown above the arrows). Red asterisks indicate the fragments are further amplified with natural stands of *P. deltoides*. (**D**) Agarose gel electrophoresis profile for fragment F_657 in females and males. M, molecular marker. B, blank control.

As the X chromosome of our sequenced male was inherited from his sequenced female parent, we could use single nucleotide polymorphisms (SNPs) to reconstruct his complete haplotypes in the region, SLR-X and SLR-Y (Fig. 1B). Interestingly, the Y haplotype includes two hemizygous fragments (which we term YHF), one long and one short, suggesting insertions into the SLR-Y, or deletions of SLR-X regions (Fig. 1B). Furthermore, the telomeres of the two haplotypes differ, with the size of the X haplotype’s telomere being 6.2 kb, and much longer (24.5 kb) for the Y haplotype (Fig. 1B). Divergence between SLR-X and SLR-Y was also higher for sequences neighboring the telomeres than in the rest of the SLR, and sequences were often unalignable (Fig. 1B, Supplementary Fig. 2). Taken together, these observations suggest that crossing over is suppressed, or greatly limited, in a small region at the SLR-end of chromosome XIX, allowing maintenance of differentiated Y and X haplotypes in a sex chromosome-like region.

We validated our haplotype reconstructions by amplifying and Sanger sequencing 13 overlapping fragments spanning the large YHF region from the sequenced male (Fig. 1C). We obtained a 41,481 bp sequence identical with the SLR-Y sequence, which confirms that no gaps exist in the SLR-Y sequence. Sequence annotation predicted 41 genes in the SLR-X and 26 in SLR-Y (Supplementary Table 2), including a cluster of 5 tandem genes encoding leucine-rich repeats (LRR) receptor-like protein kinases in both haplotypes. At least 16 genes with assigned functions were found in both the SLR-X and SLR-Y (Supplementary Table 2).

### Identification of sex determining genes

To identify the genetic factors underlying sex determination in *P. deltoides*, we performed a genome-wide association study (GWAS) based on SNPs in 49 females and 46 males (Supplementary Table 3). Genome resequencing generated a total of 1.15 Tb Illumina reads with sequence depths of at least 20× for each of the 95 sampled trees (Supplementary Table 3). With the sequenced female as the reference, we detected 435 SNPs with genotypes matching the individuals’ sexes under male heterogamety, in other words SNPs that are homozygous in all females in our samples, but heterozygous in all the males (Fig. 2A and Supplementary Table 4; we refer to these SNPs as SEMSs). Most (315) SEMSs are in three SLR genes, T-complex protein 1 subunit gamma (*TCP*), Chloride channel protein CLC-c (*CLC*), and DNA-methyltransferase 1 (*MET1*). Of the remaining 120 SEMSs, located outside the SLR, 78 are in a gene we named *FERR*, which is in the PAR of chromosome XIX (Fig. 1A), and the others are on three other chromosomes and one unplaced contig, Contig01665 (Fig. 2A). The others are probably false positives; 27 from an autosomal gene, *HEMA1*, on chromosome IX, and 15 from non-coding sequences, or genes with unknown functions (Supplementary Table 4). Previous studies of *P. balsamifera* (*24*) and *P. trichocarpa* (*24, 27*), based on using the assembled female *P. trichocarpa* sequence (*28*) as the reference genome, also found many SNPs significantly associated with sex distributed on multiple chromosomes. Using a female genome as the reference, however, is likely to cause erroneous mapping of male reads from Y-linked regions that are missing from the female genome to homologous sequences elsewhere in the genome, producing false positive SEMSs. Examination of our *P. deltoides* non-SLR SEMSs indeed revealed sequence similarity with the YHF (Supplementary Table 5). We therefore used the SLR-Y as our reference for GWAS analysis, which eliminated all the non-SLR SEMSs (Fig. 2C).

**Fig. 2.**
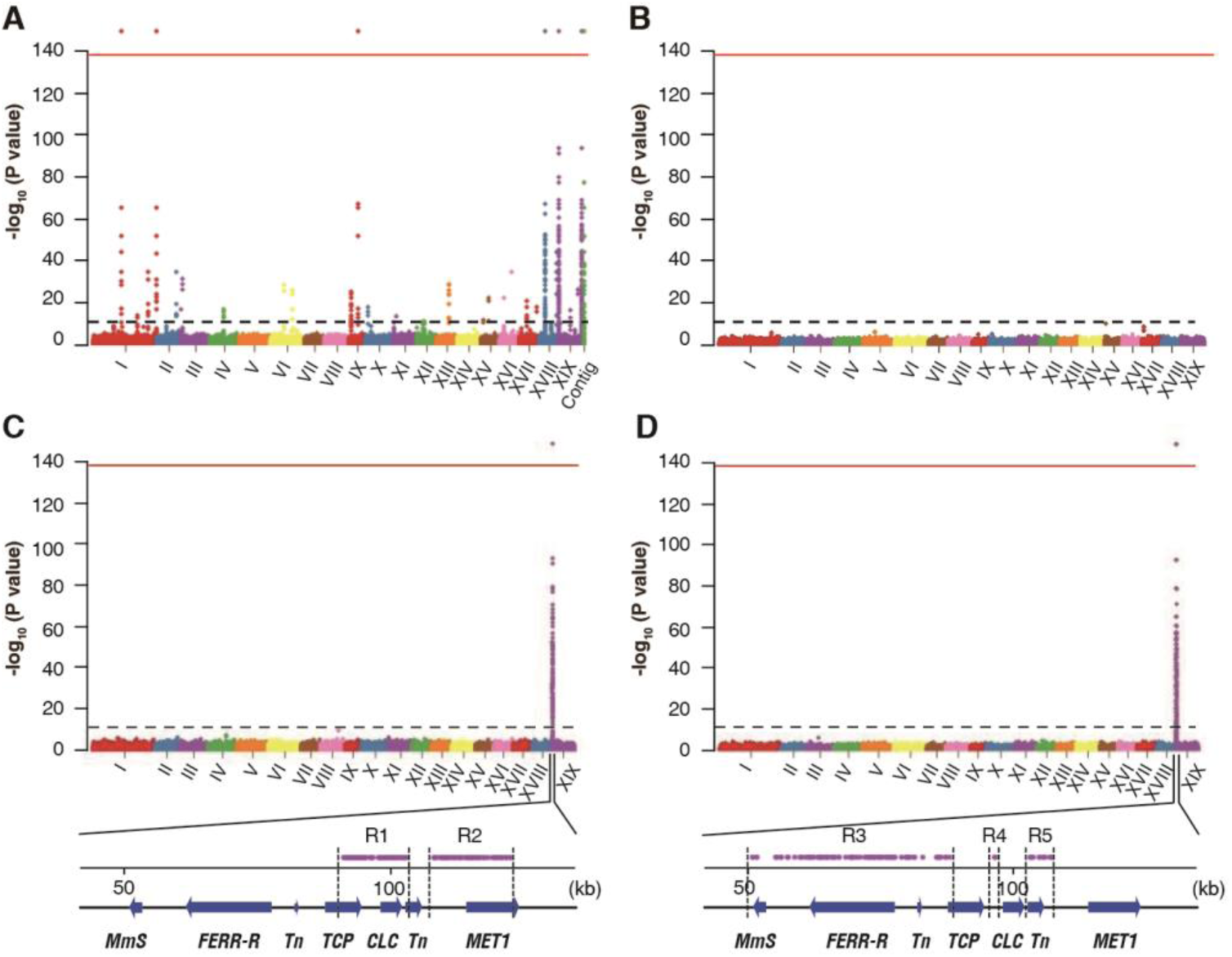
Manhattan plots of GWAS analysis on sex of *P. deltoides*. (**A-B**) Results when the female genome was used as the reference in our GWAS analyses (A shows the SNP analysis B shows the analysis of coverage). “Contig” on the *x*-axis in (A) represents the unplaced Contig01665. The *x*-axis shows the chromosome numbers, the *y*-axis shows the negative logarithm of P values, the dashed lines above the *x*-axes indicate the Bonferroni cut off 0.01, the red lines above the *x*-axes indicate the Bonferroni cut off 1e-140. (**C-D**) Results using SLR-Y as the reference genome in GWAS analyses using either SNPs (C) or coverage (D). Genes completely associated with sex are listed in the diagram of chromosome XIX below parts C and D. R1-R5 indicate regions containing variants above red lines.

As mentioned above, comparison of the SLR-X and -Y revealed two YHFs. To identify female- or male-specific hemizygous sequences, we analyzed read-coverage in the 95 *P. deltoides* GWAS samples, searching for sequences that are present in all trees of one sex, but absent in all individuals of the other sex. With the SLR-X as the reference, no female-specific hemizygous sequences were detected (Fig. 2B), but using the SLR-Y, we detected two male-specific hemizygous sequences (Fig. 2D). Both were consistent with the two YHFs shown in Fig. 1B, indicating that both of them are fully Y-linked features indicating males in *P. deltoides*, and supporting the presence of a non-recombining region. To further evaluate this result, we tested the longer YHF using PCR primers designed to amplify five separate fragments (Fig. 1C), in 20 trees of each sex from the GWAS samples; PCR amplification succeeded in all of the males but failed in all females (Fig. 1D), confirming male-specificity. The long YHF contains two non-protein-coding genes (referred to as *FERR-R* and *MmS* in Fig. 1B) and a transposable element, whereas the short YHF harbored only a transposon, presumably an insertion that is fixed in the fully Y-linked region.

### Expression of candidate sex determining genes

Poplar catkins consist of many small flowers without petals or sepals. The diminutive florets are attached to the rachis of morphologically different male or female catkins. A single male floret consists of a group of stamens inserted on a disk, while a single female floret includes a single-celled ovary seated in a cup-shaped disk (Supplementary Fig. 3). Poplars bloom in early spring before the flush of leaves, but differentiation of female and male flower primordia starts in June of the previous year (Fig. 3A). In different dioecious plants, separation of the sexes occurs at different developmental stages, and may involve different sets of genes. Longitudinal sections of flower buds (Fig. 3B and Supplementary Fig. 4) showed that *P. deltoides* male and female flower primordia are distinguishable starting from early June (T1) and early July (T3), respectively. Four stages of sexual organ abortion are recognized, stage T0 (before the initiation of stamen or carpel primordia), stage T1 (early stamen or carpel development), pre-meiosis (stage T2), and post-meiosis (stage T3) (*29*). In poplars, unisexual flower primordia form at stage T0, and thus the sex-determining genes in poplars must act at this stage to trigger the initiation of either gynoecia or androecia primordia.

**Fig. 3.**
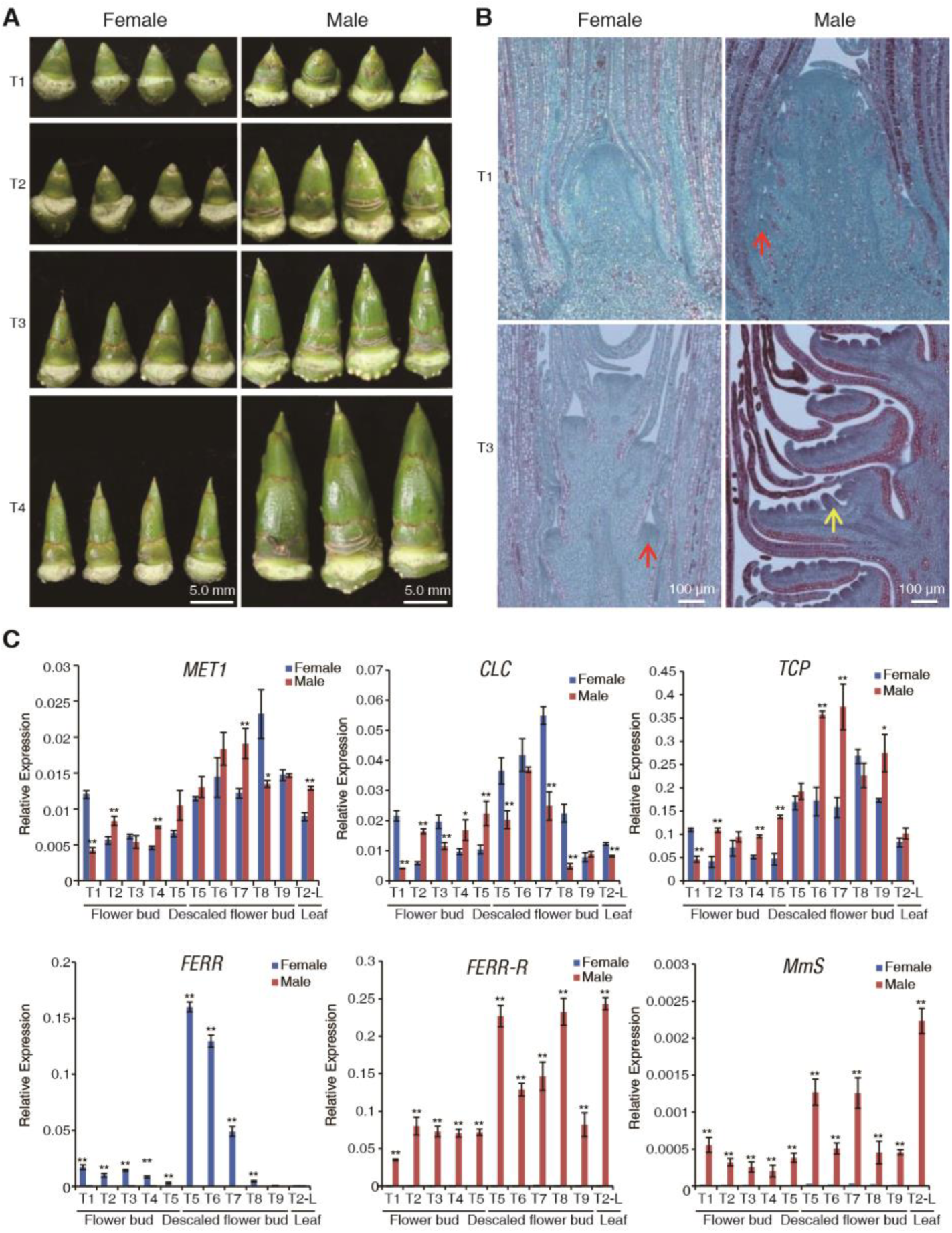
Development of flower buds and gene expression profiles. (**A**) Morphology of the male and female flower buds at developmental stages T1-T4 defined in the text. (**B**) Longitudinal sections of male and female inflorescences at T1 and T3. The red arrows point to the floret primordia, and the yellow arrow denotes the anther primordium. (**C**) Gene expression profiles. Values represents the mean ± SD from three biological replicates (*P < 0.05; **P < 0.01, Student’s t-test).

RNA-seq data detected no expression of the two Y-specific transposable elements (Fig. 1B). In contrast, *TCP, CLC, MET1, FERR-R*, and *MmS* were expressed at all sampling times examined, and qRT-PCR bioassays showed that their expression is not limited to flower tissue (Fig. 3C). No consistent difference in expression between the sexes was observed for three of these genes, *TCP, CLC*, and *MET1* (Fig. 3C), all of which are present in both the SLR-X and -Y. These three protein-coding genes cannot therefore be the sex determining genes. In contrast, the two SLR-Y hemizygous genes, *FERR-R* and *MmS* show male-specific expression (Fig. 3C), suggesting that they may control sex in *P. deltoides*.

### *FERR-R* is a femaleness suppressor that generates siRNAs suppressing *FERR* function

Sequence alignment revealed homology of the SLR-Y hemizygous *FERR-R* gene with *FERR* (located in the PAR) and *HEMA1* (an autosomal gene) (Fig. 4A). Seven homologous segments (S1, S2, S3, S4, S6, S7, and S8) were detected between the two, including all parts of *FERR* (the promoter region, 5’-UTR, exon 1, exon 2, exon 3, the first three introns, and two downstream segment). In contrast, the *FERR-R* segment with homology to *HEMA1*, named S5, corresponds only to the *HEMA1* 5’-UTR and exon 1 (Fig. 4A).

**Fig. 4.**
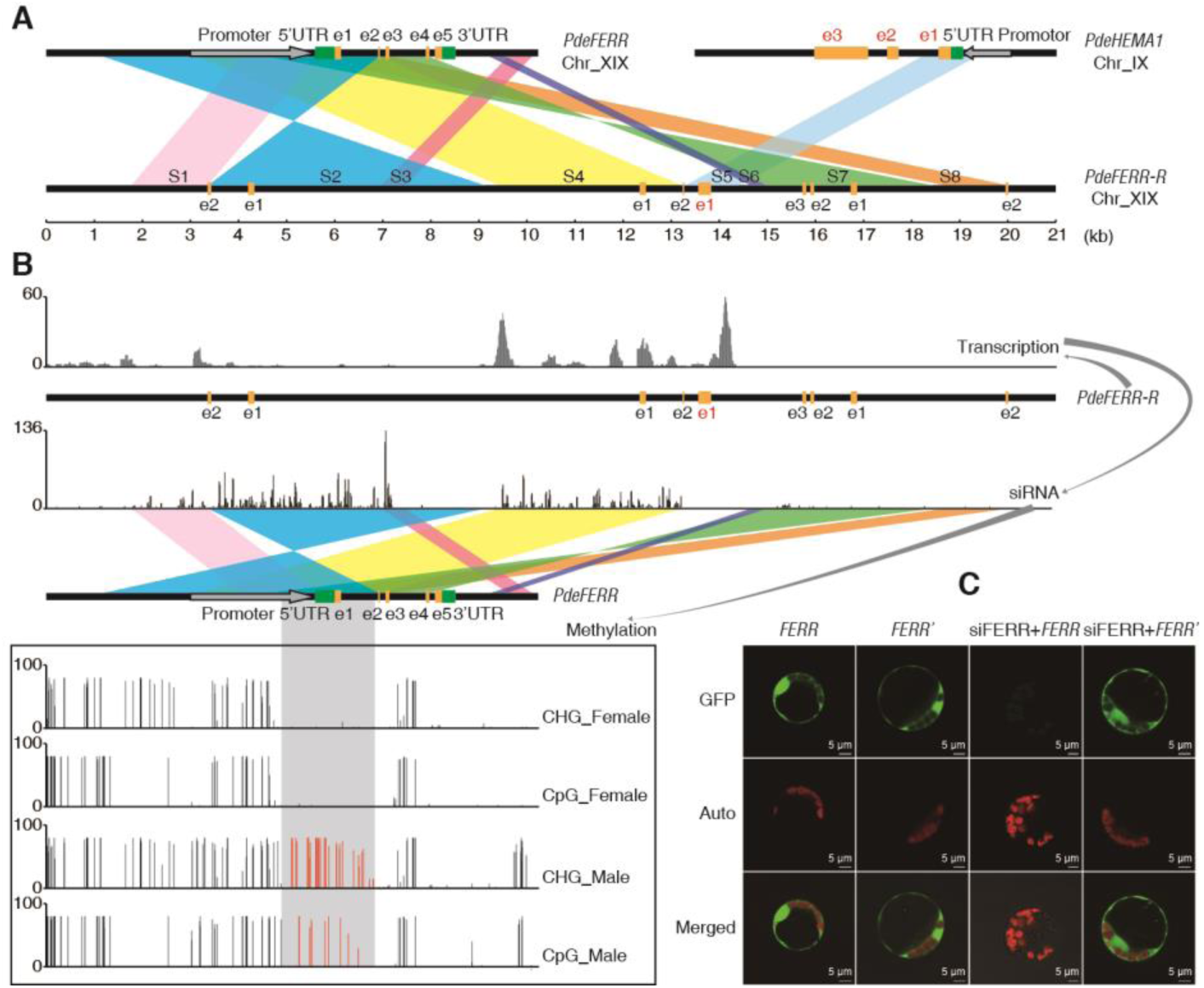
Origin of *FERR*-*R* and its function in repressing *FERR*. (**A**) Sequence homology analysis for *FERR*-*R*. e1-e5 represent the five exons (in black) of *FERR* or *HEMA1* (in red). S1-S8 indicate the duplicated segments described in the text. (**B**) Abundance of *FERR*-R transcripts and the *FERR*-*R* generated siRNAs, and the differential methylation of *FERR* in the two sexes. The gray shadow shows the region methylated only in males, and the red vertical bars indicate the methylation levels in this region. (**C**) Transient expression experiment in poplar protoplasts. siFERR is siRNA generated by *FERR*-*R. FERR’* is the siRNA resistant version of *FERR*.

Expression data from strand-specific lncRNA-Seq and small RNA-Seq revealed that *FERR-R* is transcribed into long transcripts that generate small interfering RNAs (siRNAs) (Fig. 4B). In other organisms, siRNAs have been found to guide the methylation of homologous DNA through RNA-directed DNA methylation (*30*). We found that, in *P. deltoides*, siRNAs generated by *FERR-R* could guide methylation at the promoter, 5’-UTR, exon 1, and the first intron of the *FERR* gene (Fig. 4B). Bisulfite sequencing showed that methylation of the corresponding regions in *FERR* occurred specifically in males (Fig. 4B). Male-specific DNA methylation of the promoter and the first intron in *PbaRR9*, the *P. balsamifera* locus homologous to the *P. deltoides FERR*, was also detected in *P. balsamifera* (*25*). Besides inducing siRNAs-directed DNA methylation, siRNAs produced by *FERR-R* were also found to target exon 1, exon 2 and exon 3 of *FERR*, suggesting that *FERR-R* might also trigger siRNA-guided cleavage of *FERR* transcripts. Transient expression experiment in poplar mesophyll protoplasts confirmed that cleavage indeed occurred, and involved interaction between *FERR-R* and *FERR*. Green fluorescence signals were observed in poplar protoplasts transformed with *FERR, FERR*’ (an siRNA-resistant version of *FERR*), and after co-transformation with siFERR+*FERR*’, but not after co-transformation of siFERR+*FERR* (Fig. 4C). A working model of the interaction between *FERR*-*R* and *FERR* is shown in Supplementary Fig. 5.

We propose that *FERR* is a female-specifically expressed response regulator in poplar (the “RR” in its name refers to this proposed function). Consistent with this hypothesis, and with a function in the initiation of female flower primordia and early carpel development, a qRT-PCR experiment revealed that *FERR* is expressed only during the initiation of female flower primordia and the early development of female flowers, as expected under our proposed mechanism. At developmental stage T5, *FERR* was expressed more than 50 times higher in early stage flower buds (without scales) than in those with scales, indicating higher expression in flower tissue before T5 than in later flower buds. *FERR* expression decreased to a low level at times after T8 (Fig. 3C). Overexpression of *PdeFERR* in *Arabidopsis thaliana* often yielded a phenotype of stigma exsertion, with some extreme cases of flowers with two pistils or with carpel-like sepals (Fig. 5A). In contrast, stamens were not affected. These results provide evidence that *FERR* promotes female functions, consistent with the Y-linked *FERR-R* gene suppressing its functions in *P. deltoides* males, and corresponding to the hypothesized female suppressor, or Su^F^, involved in the evolution of dioecy (*9*).

**Fig. 5.**
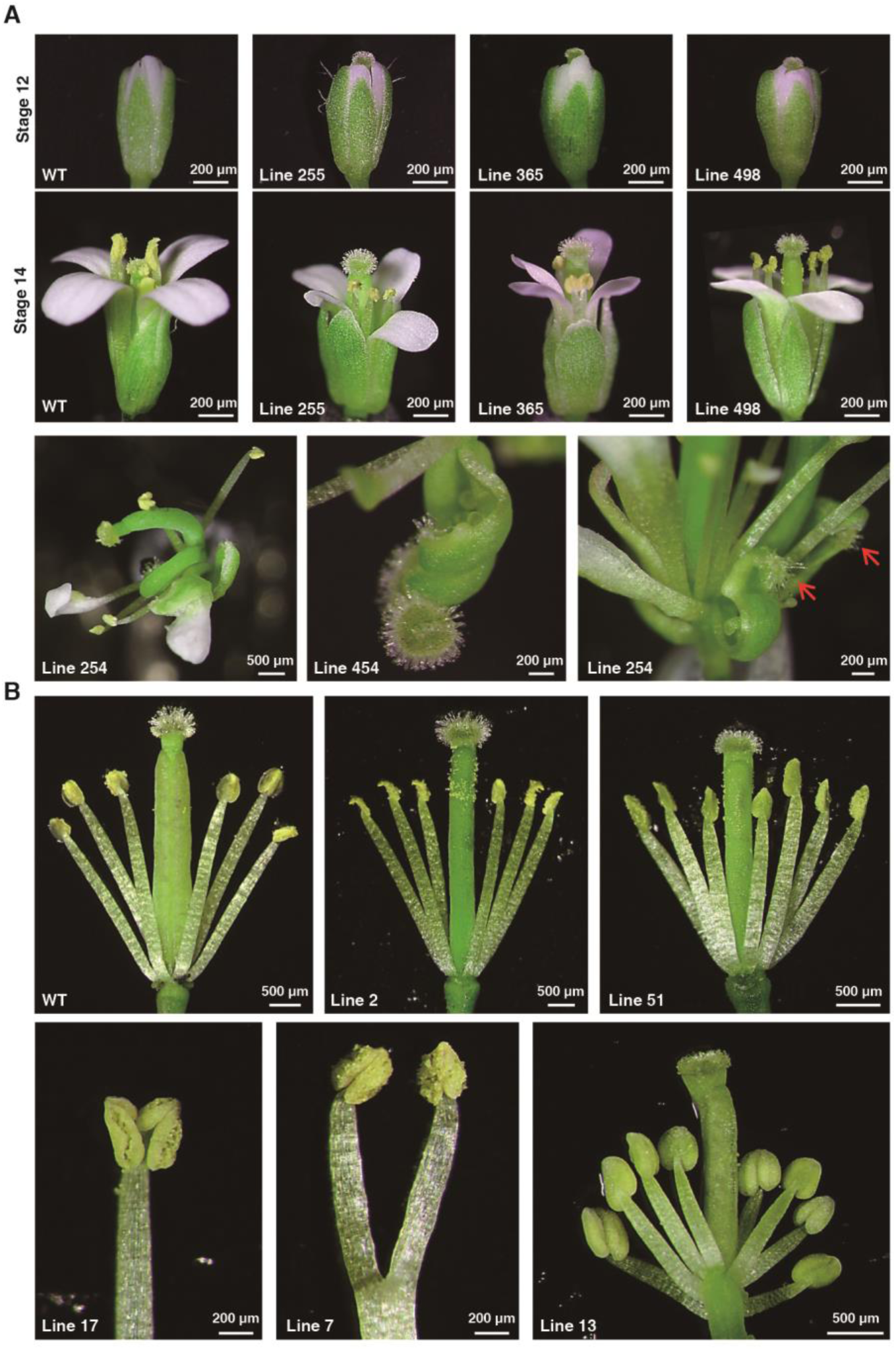
Phenotypes of transgenic *Arabidopsis.* (**A**) Phenotypes of *FERR* overexpression lines. The red arrows point to the carpel-like sepals. (**B**) Phenotypes of *MmS* overexpression lines.

### *The MmS* gene locus reduces miRNA levels and promotes maleness

The second SLR YHM gene with male-specific expression is *MmS*. We hypothesize that this gene generates miRNA molecules that promote maleness by removing transcripts that, in the non-dioecious ancestral species, reduced male functions. Bioinformatics analysis showed that multiple miRNAs are bound by *MmS* transcripts (Fig. 6A and Supplementary Table 6). Quantification by qRT-PCR revealed a continuous expression of *MmS* in male *P. deltoides* (Fig. 3C). We propose the following model for the regulatory function of *MmS* (Supplementary Fig. 6). The *MmS* transcripts bind multiple miRNAs, resulting in fewer miRNAs working on cleaving the transcripts of their target genes in males than in females. Transcripts of the target genes would thus be more abundant in males. Analysis with the tool psRNAtarget (*31*) indicated that *MmS* transcripts did not bind siRNAs produced by *FERR-R*. Therefore, *MmS* and *FERR-R* should work independently.

**Fig. 6.**
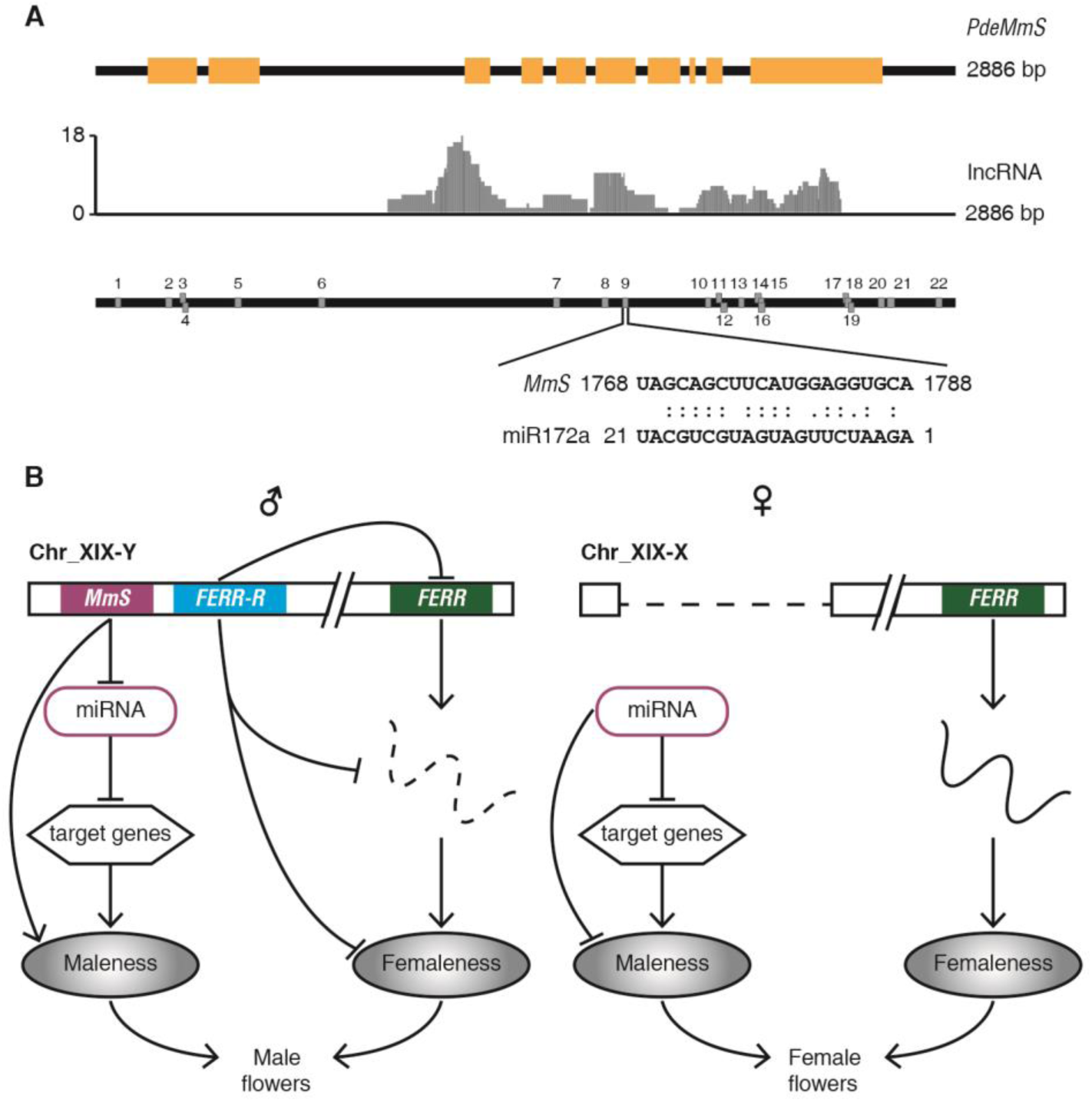
Transcription of *MmS* and the proposed model for the sex-determining genes’ action in *P. deltoides*. (**A**) Gene structure and transcription of *MmS*, and the miRNA binding sites in the *MmS* transcript. The yellow boxes indicate predicted *MmS* exons. The middle track visualizes the abundance of lncRNA. The miRNAs binding sites are numbered in the bottom track, and binding site 9 is zoomed in. (**B)** The proposed model for the sex determination genes.

Flowers of wild-type *A. thaliana* have four long and two short stamens. Overexpression of *PdeMmS* in *Arabidopsis* affected androecium phenotypes, commonly resulting in flowers with six long stamens, seven or even occasionally 8 stamens, stamens bearing two anthers, or branched stamens (Fig. 5B). In contrast to stamen, pistil was not affected by overexpression of *PdeMmS*. These phenotypes are consistent with *MmS* being a maleness promotor, working independently of *FERR-R*.

### Sex determination in *P. davidiana*

*P. deltoides* belongs to subgenus *Aigeiros* in genus *Populus*. To test whether the YHFs is conserved in other poplars, we sequenced the genome of *P. davidiana*, which belongs to subgenus *Leuce*, an earlier-branching section of *Populus* than *Aigeiros* (32). First, a male *P. davidiana* was sequenced and assembled, and used as the reference genome. Subsequent genomic re-sequencing was applied to the GWAS samples of *P. davidiana*. Coverage analysis detected a YHF of 126 kb on contig ctg345 (Supplementary Fig. 7), which again includes two of the same genes as the larger *P. deltoides* YHF, *FERR-R* (named *PdaFERR-R*) and *MmS*. Twelve duplicated segments were detected between *PdaFERR-R* and the *PdaFERR* gene, the putative source of the duplication that created the YHF. However, the duplicated exons differ between *P. deltoides* and *P. davidiana* (Fig. 4A; Supplementary Fig. 7). In *P. davidiana* only exon 1 of *FERR* is duplicated and no *HEMA1* exons are duplicated into *FERR-R*. Our results therefore suggest that separation of sexes in these two poplar species occurred independently, involving a similar mechanism and the same genes, but with different duplicated segments.

## Discussion

Many genes probably function in the development of sex dimorphisms of poplars, but *FERR-R* and *MmS* appear to be the upstream genes determining sex in *P. deltoides*. We propose that, in *P. deltoides* XX females, *FERR* function is active due to the absence of the *FERR-R* gene (Supplementary Fig. 5), whereas *MmS* in *P. deltoides* XY males appears to function through removal of mRNA targets via a miRNA “sponge” mechanism (Supplementary Fig. 6). Under the two-gene model outlined above (Fig. 6B), the mechanism revealed in this study can explain the evolution of separate sexes from a hermaphroditic ancestor (Supplementary Fig. 8).

A distinctive feature of the many Y chromosomes, including the mammalian, bird and Drosophila Ys, is absence of crossing over, sometimes followed by loss of large numbers of genes present on the X (and on the Y ancestor) by the process known as genetic degeneration (*33, 34*). The *P. deltoides* sex-determining genes also appear to be in a non-recombining region, but it is physically much smaller than the animal cases just mentioned, and even than the small regions in kiwifruit (*35*) and asparagus (*7*). Our results suggest why the region that carries the two *P. deltoides* sex-determining genes does not recombine, and also suggest that there are two different routes by which recombination is prevented between plant Y- and X-linked regions.

Epigenetic regulation of reproductive genes is common in sex determination mechanisms in plants and animals (*36*), including in the sex-determining mechanism by which monoecious plants develop male or female flowers in differing developmental contexts, rather than under control of different plant genotypes (*37*). Specifically, miRNAs and lncRNAs have been found to function in sexual reproduction (*38, 39*). Our study shows that, in poplars, *FERR-R* function involves both RNAi and DNA methylation processes, with *MmS* being the first reported miRNA sponge gene triggering sex separation in dioecious plants. Since many miRNA families are evolutionarily conserved across all major lineages of plants (*40*), the *MmS* gene may be important for the formation of male organs in plants other than poplars.

Poplars dominant the landscape in many regions of the world, and provide an important commercial source of fiber and fuel (*41*). The independent functions of sex determining genes enables us to block gynoecia and androecia development in female and male poplars respectively via gene editing. This study provides genes for modification to reduce the energy consumption in sexual reproduction, and to resolve the heavy air pollutions caused by seed-hairs from the female and pollens from the male poplars.

## Acknowlegments

This work is supported by the National Key Research 300 Project of China (2016YFD0600101), and grants from Natural Science Foundation of China (31561123001) and NSF1542599.

## Author Contributions

T.Y. designed the experiments. H.W., Y.C., X.L., J.H., J.L., and X.D. performed the experiments. L.X. and S.W. analyzed the data. M.X. and T. Y. maintained the poplar collections and the mapping population. T.Y., L.X., H.W., J.H., Y.C., X.L., M.O., and J.Q.L. drafted the manuscript. D.C. led the interpretation of the theoretical perspectives and critically reformulated the manuscript. All authors participated in data interpretation and approved the final manuscript.

## Materials and Methods

### Plant materials

To map the sex locus, an intraspecific F_2_ population of *P. deltoides* was established in 2012 by crossing two randomly selected siblings from an F_1_ full-sib family. A total of 1,077 offspring (550 females and 527 males) of the F_2_ population were planted (6×6m spacing) and maintained at Sihong Forest Farm in Jiangsu, China. To obtain the reference genomes for female and male *P. deltoides*, we sequenced the maternal parent and a randomly selected male progeny from the F_2_ mapping pedigree. For genome-wide association study (GWAS), we sequenced the genomes of 49 unrelated females and 46 unrelated males (referenced as GWAS samples of *P. deltoides*), which were selected from the *P. deltoides* germplasm collections maintained at Sihong Forest Farm. The *P. deltoides* germplasms plantation (6×6m spacing) was established with 12 ramets for each clone following a completely random block design in 1998.

To characterize the developmental processes of flower buds, we collected the female and male flower buds from a female and a male tree in *P. deltoides* germplasms at nine different times: T1 (June 3), T2 (June 18), T3 (July 3), T4 (July 18), T5 (August 3), T6 (August 18), T7 (September 3), T8 (December 1), and T9 (January 15). To quantify the expression levels of genes, flower buds at the corresponding times were separately collected from three ramets of the female and male *P. deltoides* for biological replications. In the early developmental stages, it is difficult to harvest enough tissue for molecular experiments, and thus the flower buds with scales were used for T1-T5. For T5-T9, the descaled flower buds were used in molecular experiments. Since flower tissue only accounts for a small portion of the flower bud, we measured expression by using both the scaled and descaled flowers at T5 for comparison purposes.

Apart from *P. deltoides*, we also explored the sex determination in *P. davidiana*, a more primitive species belonging to the section of *Leuce* in genus of *Populus* (*32*). To study the sex determination in *P. davidiana*, leaf samples were collected from a natural population of *P. davidiana* along Datong River in Qinghai, China in spring 2019. GWAS samples of *P. davidiana* included 49 females and 47 males.

### Sequencing female and male *P. deltoides* genomes and mapping the sex locus

Young leaves of the maternal parent and a randomly selected male progeny were collected for DNA extraction and sequencing libraries construction. For PacBio sequencing, high molecular weight DNA was extracted using the cetyltrimethylammonium bromide method (*42*). g-TUBE (Covaris, USA) was used to shear DNAs of the female and male separately into fragments with an average size of 20 kb. The PacBio SMRT libraries were constructed using sheared DNAs, then sequenced following standard protocols (PacBio, USA). The initial assemblies of PacBio reads of the female were generated by using Wtdbg v2.5 (*43*), Falcon v0.3.0 (*44*), and Canu v1.8 softwares (*45*). Quickmerge was applied to combine the assemblies mainly based on Wtdbg assembly (*43*). The resulted sequences were further polished using Illumina genomic resequencing reads with Pilon v1.23 software (*46*). Hi-C libraries were constructed for the female parent following the Proximo™ Hi-C plant protocol (Phase Genomics, USA) and applied for sequencing on an Illumina HiSeq X platform (Illumina, USA). HiC-Pro v2.10.0 (*47*) was used to evaluate the quality of HiC data, and reads of valid interaction pairs were applied for genome scaffolding using Lachesis v20171221 (*48*). BUSCO v3.0.2 (*49*) was used to evaluate the completeness of the genome assembly.

To map the sex locus, complete genetic maps were built for the female and male parents with 94 randomly selected offspring by using AFLP markers following the two-way pseudo-testcross strategy (*50*). Generation of AFLP markers was conducted following the description in Yin *et al.* (*51*). With the established genetic maps, whole genome scan for sex locus was performed to locate its general position. Referring to the poplar consensus map (*5*), we developed simple sequence repeat (SSR) markers in the vicinity region of the sex locus. The designed SSRs were then genotyped and mapped on the AFLP maps to saturate the target chromosomal region. In the mapped SSRs, we further selected six closely sex-linked SSRs to conduct fine local mapping. The selected SSRs were genotyped with 1,077 flowered progenies to confine the sex locus in a more precise interval. To ensure the accuracy of sex phenotyping, the sex of each tree was separately recorded by three teams in two rounds of observation.

### Genome annotation

A *de novo* repeat library was constructed using LTR FINDER v1.05 (*52*), RepeatScout v1.0.5 (*53*), and PILER-DF v2.4 (*54*). The resulted repeat elements were categorized using PASTEC lassifier v1.0 (*55*) and combined with Repbase (*56*), which were further imported to RepeatMasker (version 4.07) (*57*) to identify and cluster repetitive elements. Sequences with more than ten monomers simple repeats ‘CCCTAAA’ were identified as telomeres.

Evidence of multiple resources was used to predict protein-coding genes in the genome. RNA-Seq data were mapped to the reference genome using HISAT2 v2.0.5 (*58*) and assembled into transcripts using Stringtie v1.3.6. (*59*). The resulted transcripts were screened using TransDecoder v5.0.2 (*60*) and GeneMarkS-T v5.1 (*61*) for protein-coding genes. PASA v2.0.2 (*62*) was used to predict gene structures of the transcripts. The *ab initio* prediction was performed using Genscan (*63*), Augustus v3.2.3 (*64*), and SNAP v2013-11-29 (*65*). GeMoMa v1.5.3 (*66*) was applied for homology-based predictions. All the annotations were integrated using EVM v1.1.1 (*67*). Infenal v1.1.2 (*68*) was used to predict rRNAs using Rfam as reference. The tRNA genes were predicted using tRNAscan-SE v1.3.1 (*69*). Pseudogenes were predicted using GeneWise v2.4.1 (*70*) after the filtering of protein-coding genes using GenBlastA v1.0.4 (*71*). The functions of genes were annotated through similarity search of databases including NR, KOG, KEGG, and TrEMBL.

### Constructing the SDR haplotypes

To compare the sequences in SDR between the X and Y chromatids, we reconstructed the haplotypes of SDR-X and SDR-Y as follows: Canu software (*45*) was applied to assemble the PacBio reads of the sequenced male. The purge_haplotigs tool v1.0.3 (*72*) was used to screen primary contigs and allelic secondary contigs. With the sequenced female as reference, both the primary and the secondary contigs were mapped to the reference genome. The contigs located in SDR were assigned to X and Y haplotypes based on SNPs between contigs and the reference genome.

To validate the reconstruction for the Y-specific region, sequence-specific primers were designed according to SDR-Y, and these primers were used to amplify DNAs extracted from the sequenced male. All PCR reactions were performed with PrimeSTAR® GXL Premix (Takara, China) according to the user manual. Details of the primers and PCR conditions were listed in Extended Data Table 1. The amplification products of each reaction were purified using the AxyPrep DNA Gel Extraction Kit (Axygen Scientific, USA), cloned into pEASY-Blunt vector (Transgen Biotech, China), and then sequenced on the Sanger sequencing platform ABI 3730xl (Applied Biosystems, USA). The obtained sequences were assembled into an integrated contig and aligned to SDR-Y to evaluate the reliability of the reconstruction for the Y-specific region.

The conservatism of the Y-specific region was further confirmed by PCR amplification with Yspecific primer pairs of 651F/R, 655F/R, 657F/R, 661F/R, and 712F/R (Extended Data Table 1) against DNAs extracted from 20 male and 20 female *P. deltoides*, which were randomly selected from the GWAS samples. The amplification products were examined by electrophoresis on 1% agarose.

### GWAS analysis on sex determination

Genomic resequencing was performed for the GWAS samples using Illumina platform. After removing low-quality reads, the Trimmonatic v0.36 (*73*) was used to trim the adaptor sequences from read ends, and BWA v0.7.17 (*74*) was used to map the Illumina reads onto the reference genomes. Two versions of references were applied for GWAS. The consensus sequence of the female genome was selected in the first version, whereas the second version SDR-X was substituted by SDR-Y. Freebayes v0.9.10 (*75*) was applied to call variants including SNPs, short Indels, and MNPs. For the sake of brevity, all these variants were referred to as SNPs in our analyses. The variants (SNPs) were screened at population level using plink v1.9 (*76*) with settings “--maf 0.05 --geno 0.1” and converted into gene dose file using qctool v2.0.1. GEMMA v0.98.1 software (*77*) was used to perform SNP-based GWAS (snp-GWAS) between SNPs and sex phenotypes.

In read-coverage based GWAS (rb-GWAS), 100-bp windows were generated across the whole genome, the read depths were calculated using bedtools v2.27.1 (*78*). Windows with maximum coverage of 0, 1-2, and >=3 were assigned with doses of 0, 1 and 2. The windows associated with sex phenotypes were identified using GEMMA following the description as in snp-GWAS.

### Characterizing flower development and quantifying genes expression

Flower buds at different times (T1-T4) were collected and fixed using formalin-acetic acid. The samples were further dehydrated in a graded concentration of ethanol and embedded in paraplast (*79*). Serial sections were prepared by employing a Leica microtome. The sections were then mounted on microscope slides for staining with 1% safranin and examined using Carl Zeiss Imager M2 microscope (Zeiss, Germany). RNA prep Pure Plant Kit (Tiangen, China) was used to extract RNAs from flower buds and leaf tissues. TransScript One-Step gDNA Removal and cDNA Synthesis SuperMix (TransGen Biotec, China) were used to reversely transcribe RNA to cDNA. cDNAs transcribed by Oligo dT primer were used to analyze the relative expression of coding genes, and those transcribed with random primers were used to analyze the relative expression of lncRNAs. AceQ qPCR SYBR Green Master Mix (Vazyme, China) was used to perform Quantitative real-time PCR (qRTPCR) on a 7500 Fast Real-Time PCR System (Applied Biosystems, USA). In each 20 μl reaction volume, 100 ng of cDNA was used as templates. The PCR parameters used were as follows: 95°C for 3 min, 40 cycles of 95 °C for 15 seconds, 60 °C for 15 seconds, and 72 °C for 30 seconds. Gene -specific primers were designed for *MET1, CLC, TCP, FERR, FERR-R*, and *MmS* (Extended Data Table 1). *Populus UBIQUITIN* (*PtUBQ*) gene was selected as an internal reference (*80*). The relative expression levels were calculated using the 2^-ΔΔCT^ method (*81*). The mean values and standard errors were calculated based on expression data of three biological replicates.

### Quantifying the digital expression of genes and analyzing the interaction between *FERR-R* and *FERR*

To quantifying the digital expression of genes and to explore gene regulation patterns, Illumina sequencing experiments (Illumina NovaSeq 6000, USA) were performed at levels of mRNA, lncRNA, small RNA and DNA methylation. RNA-Seq was performed to quantify the expression of the four protein-coding genes (*MET1, CLC, TCP, FERR*); whereas strand-specific lncRNASeq was applied for measuring the transcription levels of the two non-protein-coding genes (*FERR-R, MmS*), as lncRNA-Seq worked for transcripts with and without polyA tails. The samples applied for sequencing were listed in Extended Data Table 7. Libraries were constructed and sequenced following instructions from manufacturers of biochemical kits and sequencing equipment.

### Quantitative analysis of Illumina reads

Before applying to the corresponding analysis pipeline, low-quality Illumina reads were removed, and adaptor sequences were trimmed using Trimmonatic v0.36 (*73*). In data analysis of RNA-Seq, lncRNA-Seq, and sRNA-Seq, the rRNA/tRNA contaminations were also removed. STAR v2.5.3a (*82*) was used to map the RNA-Seq and lncRNA-Seq reads onto the reference, and DEseq2 (*83*) was applied for differential expression analysis.

### Experimental verification of interaction between *FERR-R* and *FERR*

Isolation of *Populus* mesophyll protoplasts was performed using the PEG-Mediated plant protoplast transformation kit (Shanghai Maokang Biotechnology, China) with 3% (w/v) cellulose R10 (Yakult Pharmaceutical, Japan) and 0.8% (w/v) macerozyme R10 (Yakult Pharmaceutical, Japan). The middle section of expanded *Populus* leaves (micropropagated *Populus* clone ‘Nanlin 895’) were cut into 0.5-1 mm fine strips, digested for 30 min in the dark using a desiccator for Vacuum infiltrate, and succeeded with digestion in the dark for 5 h without shaking (*84*). The protoplasts were harvested by filtering through a 70 μm pore nylon cloth and then suspended in the transfection buffer.

The sequence of one siRNA generated from *FERR-R* locus was designed onto the backbone of AtMIR172a, driven by CaMV 35S promoter. The artificial miRNA generated in the construction mimics the siRNA of 21nt from *FERR-R*. The artificial miRNA precursor sequence was synthesized on a Dr. Oligo384 (Biolytic, USA) and constructed into the vector p2GWF7 using Gateway technology (*85*). The full length of the *FERR* was amplified by using PrimeSTAR Max (TaKaRa, China) from cDNA of female flower bud. Site-directed synonymous mutagenesis of *FERR* was performed using the single-tube ‘megaprimer’ PCR method (*86*) to generate the resistant version of *FERR* (referred to as *FERR*’). *FERR*’ transcript was synthesized with synonymous substitutions in the complementary sequences of artificial siRNA, which was also included in the experiment to test the functional specificity between *FERR-R* and *FERR*. Pro35S::PtFERR-GFP and Pro35S::FERR’-GFP were co-transfected with Pro35S::siFERR into *Populus* protoplasts. GFP fluorescence was captured using CarlZeiss LSM710 confocal microscope (Zeiss, Germany).

### Annotation of miRNA sponges and the interacting miRNAs

Transcripts of lncRNA were assembled using Trinity V2.6.6 (*60*) with the parameter settings for strand-specific reads. The obtained sequences were applied to search NR and Sprot databases to identify homology protein-coding or non-coding transcripts. The interacting pairs of non-coding transcripts and known miRNAs in *Populus* were predicted using psRNATarget (*31*) with an increased limit of mismatch score to 6, which ensured the binding of miRNAs to the lncRNA transcripts without cleaving of the transcripts at the binding sites (*87*).

### Confirming the functions of *FERR-R* and *MmS* in transformed *Arabidopsis*

Binary vector p2301-35Splus was used in the transgenic experiments. The vector was created by sequentially cloning of *Hind*III-*Sma*I fragment and *EcoR*I-*Sac*I fragment of pBI121 (AF485783.1) into pCAMBIA2301 (AF234316). The genomic DNAs of *FERR* and *MmS*, together with CDS of *FERR* were separately cloned into p2301-35Splus. *Arabidopsis* ecotype Columbia-0 (Col-0) was grown under white LED light (Philips, Netherlands) with 16h-light and 8h-dark cycles at 19-23 °C until transformation. The binary construct was introduced into *Agrobacterium tumefaciens* strain GV3101 (pMP90) using a freezing method. *Arabidopsis* wildtype plants were transformed using the floral dip method (*88*). Screening of transgenic plants was processed on 1/2 MS media containing 50 mg/mL kanamycin, and kanamycin-resistant transgenic seedlings were further confirmed by GUS staining. Wild type Col-0 and transgenic *Arabidopsis* plants were grown in a growth chamber at 23 °C/15 °C day/night temperatures under a 16h/8h light/dark cycle. Fresh flowers at different developmental stages were conducted for microscopic observation with an Olympus SZX10 (Olympus, Japan) when plants were 7 weeks old. Floral stages were defined according to Smyth *et al.* (*89*). For each stage, five flowers were collected from the main inflorescence of the same plant.

### Study the sex determination in *P. davidiana*

To explore the sex determination in *P. davidiana*, we sequenced the genome of a male tree using PacBio Sequel II (PacBio, USA) and assembled the genome following the same pipeline as that for *P. deltoides*. We then conducted genome resequencing for the GWAS samples of *P. davidiana* and detected the YHF in males of *P. davidiana* by using rb-GWAS analysis. Sequence annotation and evolutionary analysis for duplicated genes were performed following the same pipelines as that of *P. deltoides*.

## REFERENCES

1 Ming, R., Bendahmane, A. & Renner, S. S. Sex chromosomes in land plants. Annu. Rev. Plant Biol. 62, 485–514 (2011).

2 Renner, S. S. The relative and absolute frequencies of angiosperm sexual systems: dioecy, monoecy, gynodioecy, and an updated online database. Am. J. Bot. 101, 1588–1596 (2014).

3 Pannell, J. R. Plant sex determination. Curr. Biol. 27, R191–R197 (2017).

4 Liu, Z. et al. A primitive Y chromosome in papaya marks incipient sex chromosome evolution. Nature 427, 348 (2004).

5 Yin, T. et al. Genome structure and emerging evidence of an incipient sex chromosome in Populus. Genome Res. 18, 422–430 (2008).

6 Akagi, T., Henry, I. M., Tao, R. & Comai, L. A Y-chromosome-encoded small RNA acts as a sex determinant in persimmons. Science 346, 646–650 (2014).

7 Harkess, A. et al. The asparagus genome sheds light on the origin and evolution of a young Y chromosome. Nat. Commun. 8, 1279 (2017).

8 Westergaard, M. in Advances in Genetics. Vol. 9 217–281 (Elsevier, 1958).

9 Charlesworth, B. & Charlesworth, D. A model for the evolution of dioecy and gynodioecy. Am. Nat. 112, 975–997 (1978).

10 Akagi, T. et al. A Y-encoded suppressor of feminization arose via lineage-specific duplication of a cytokinin response regulator in kiwifruit. Plant Cell 30, 780–795 (2018).

11 Akagi, T. et al. Two Y-chromosome-encoded genes determine sex in kiwifruit. Nat. Plants 5, 801–809 (2019).

12 Dellaporta, S. L. & Calderon-Urrea, A. The sex determination process in maize. Science 266, 1501–1505 (1994).

13 Boualem, A. et al. A cucurbit androecy gene reveals how unisexual flowers develop and dioecy emerges. Science 350, 688–691 (2015).

14 Akagi, T. & Charlesworth, D. Pleiotropic effects of sex-determining genes in the evolution of dioecy in two plant species. Proc. R. Soc. B 286, 20191805 (2019).

15 Markussen, T., Pakull, B. & Fladung, M. Positioning of sex-correlated markers for Populus in a AFLP-and SSR-Marker based genetic map of *Populus tremula*×*tremuloides*. Silvae Genetica 56, 180–184 (2007).

16 Gaudet, M. et al. Genetic linkage maps of *Populus nigra* L. including AFLPs, SSRs, SNPs, and sex trait. Tree Genet. Genomes 4, 25–36 (2008).

17 Pakull, B. et al. Genetic linkage mapping in aspen (*Populus tremula* L. and *Populus tremuloides* Michx.). Tree Genet. Genomes 5, 505–515 (2009).

18 Paolucci, I. et al. Genetic linkage maps of *Populus alba* L. and comparative mapping analysis of sex determination across *Populus* species. Tree Genet. Genomes 6, 863–875 (2010).

19 Pakull, B. et al. Genetic mapping of linkage group XIX and identification of sex-linked SSR markers in a *Populus tremula× Populus tremuloides* cross. Can. J. Forest Res. 41, 245–253 (2011).

20 Tuskan, G. A. et al. The obscure events contributing to the evolution of an incipient sex chromosome in *Populus*: a retrospective working hypothesis. Tree Genet. Genomes 8, 559–571 (2012).

21 Kersten, B. et al. The sex-linked region in *Populus tremuloides* Turesson 141 corresponds to a pericentromeric region of about two million base pairs on *P. trichocarpa* chromosome 19. Plant Biology 16, 411–418 (2014).

22 Song, Y. et al. Sexual dimorphic floral development in dioecious plants revealed by transcriptome, phytohormone, and DNA methylation analysis in *Populus tomentosa*. Plant Mol. Biol. 83, 559–576 (2013).

23 Song, Y., Tian, M., Ci, D. & Zhang, D. Methylation of microRNA genes regulates gene expression in bisexual flower development in andromonoecious poplar. J. Exp. Bot. 66, 1891–1905 (2015).

24 Geraldes, A. et al. Recent Y chromosome divergence despite ancient origin of dioecy in poplars (*Populus*). Mol. Ecol. 24, 3243–3256 (2015).

25 Bräutigam, K. et al. Sexual epigenetics: gender-specific methylation of a gene in the sex determining region of *Populus balsamifera*. Sci. Rep. 7, 45388 (2017).

26 Sanderson, B. J., Wang, L., Tiffin, P., Wu, Z. & Olson, M. S. Sex-biased gene expression in flowers, but not leaves, reveals secondary sexual dimorphism in *Populus balsamifera*. New Phytol. 221, 527–539 (2019).

27 McKown, A. D. et al. Sexual homomorphism in dioecious trees: extensive tests fail to detect sexual dimorphism in *Populus*. Sci. Rep. 7, 1831 (2017).

28 Tuskan, G. A. et al. The genome of black cottonwood, *Populus trichocarpa* (Torr. & Gray). Science 313, 1596–1604 (2006).

29 Diggle, P. K. et al. Multiple developmental processes underlie sex differentiation in angiosperms. Trends Genet. 27, 368–376 (2011).

30 Matzke, M. A. & Mosher, R. A. RNA-directed DNA methylation: an epigenetic pathway of increasing complexity. Nat. Rev. Genet. 15, 394–408 (2014).

31 Dai, X. & Zhao, P. X. psRNATarget: a plant small RNA target analysis server. Nucleic Acids Res. 39, W155–W159 (2011).

32 Wang, M. et al. Phylogenomics of the genus *Populus* reveals extensive interspecific gene flow and balancing selection. New Phytol. 225, 1370–1382 (2020).

33 Charlesworth, B. & Charlesworth, D. The degeneration of Y chromosomes Phil. Trans. R. Soc. Lond. B. 355 1563–1572 (2000).

34 Charlesworth, D., Charlesworth, B. & Marais, G. Steps in the evolution of heteromorphic sex chromosomes. Heredity 95, 118–128 (2005).

35 Pilkington, S. M. et al. Genetic and cytological analyses reveal the recombination landscape of a partially differentiated plant sex chromosome in kiwifruit. BMC plant biol. 19(1), 172 (2019).

36 Piferrer, F. Epigenetics of sex determination and gonadogenesis. Dev. Dyn. 242, 360–370 (2013).

37 Parkinson, S. E., Gross, S. M. & Hollick, J. B Maize sex determination and abaxial leaf fates are canalized by a factor that maintains repressed epigenetic states. Dev. Biol. 308, 462–473 (2007).

38 Chuck, G., Meeley, R., Irish, E., Sakai, H. & Hake, S. The maize *tasselseed4* microRNA controls sex determination and meristem cell fate by targeting *Tasselseed6/indeterminate spikelet1*. Nat. Genet. 39, 1517–1521 (2007).

39 Golicz, A. A., Bhalla, P. L. & Singh, M. B. lncRNAs in plant and animal sexual reproduction. Trends Plant Sci. 23, 195–205 (2018).

40 Jones-Rhoades, M. W., Bartel, D. P. & Bartel, B. MicroRNAs and their regulatory roles in plants. Annu. Rev. Plant Biol. 57, 19–53 (2006).

41 FAO. Poplars and Other Fast - Growing Trees –Renewable Resources for Future Green Economies. Synthesis of Country Progress Reports. 25th Session of the International Poplar Commission, Berlin, Federal Republic of Germany, 13-16 September 2016. Working Paper IPC/15. Forestry Policy and Resources Division, FAO, Rome (2016).

## REFERENCES

42. Murray, M. & Thompson, W. F. Rapid isolation of high molecular weight plant DNA. Nucleic Acids Res. 8, 4321–4326 (1980).

43. Ruan, J. & Li, H. Fast and accurate long-read assembly with wtdbg2. Nat. Methods, doi.org/10.1038/s41592-019-0669-3530972 (2019).

44. Chin, C.-S. et al. Phased diploid genome assembly with single-molecule real-time sequencing. Nat. Methods 13, 1050 (2016).

45. Koren, S., Walenz, B. P., Berlin, K., Miller, J. R. & Phillippy, A. M. Canu: scalable and accurate long-read assembly via adaptive *k*-mer weighting and repeat separation. Genome Res. 27, 722–736 (2017).

46. Walker, B. J. et al. Pilon: an integrated tool for comprehensive microbial variant detection and genome assembly improvement. PLoS One 9 (2014).

47. Servant, N. et al. HiC-Pro: an optimized and flexible pipeline for Hi-C data processing. Genome Biol. 16, 259–259 (2015).

48. Burton, J. N. et al. Chromosome-scale scaffolding of *de novo* genome assemblies based on chromatin interactions. Nat. Biotechnol. 31, 1119–1125 (2013).

49. Simão, F. A., Waterhouse, R. M., Ioannidis, P., Kriventseva, E. V. & Zdobnov, E. M. BUSCO: assessing genome assembly and annotation completeness with single-copy orthologs. Bioinformatics 31, 3210–3212 (2015).

50. Grattapaglia, D. & Sederoff, R. Genetic linkage maps of *Eucalyptus grandis* and *Eucalyptus urophylla* using a pseudo-testcross: mapping strategy and RAPD markers. Genetics 137, 1121–1137 (1994).

51. Yin, T-M., Wang, X-R., Andersson, B. & Lerceteau-Köhler, E. Nearly complete genetic maps of *Pinus sylvestris* L. (Scots pine) constructed by AFLP marker analysis in a full-sib family. Theor. Appl. Genet. 106, 1075–1083 (2003).

52. Xu, Z. & Wang, H. LTR_FINDER: an efficient tool for the prediction of full-length LTR retrotransposons. Nucleic Acids Res. 35, 265–268 (2007).

53. Price, A. L., Jones, N. C. & Pevzner, P. A. *De novo* identification of repeat families in large genomes. Bioinformatics 21, i351–i358 (2005).

54. Edgar, R. C. & Myers, E. W. PILER: identification and classification of genomic repeats. Bioinformatics 21, i152–i158 (2005).

55. Hoede, C. et al. PASTEC: an automatic transposable element classification tool. PloS one 9, e91929 (2014).

56. Bao, W., Kojima, K. K. & Kohany, O. Repbase Update, a database of repetitive elements in eukaryotic genomes. Mob. DNA 6, 11 (2015).

57. Smit, A., Hubley, R. & Green, P. RepeatMasker Open-3.0. (1996).

58. Kim, D., Langmead, B. & Salzberg, S. L. HISAT: a fast spliced aligner with low memory requirements. Nat. Methods 12, 357–360 (2015).

59. Pertea, M. et al. StringTie enables improved reconstruction of a transcriptome from RNA-seq reads. Nat. Biotechnol. 33, 290–295 (2015).

60. Haas, B. J. et al. De novo transcript sequence reconstruction from RNA-seq using the Trinity platform for reference generation and analysis. Nat. Protoc. 8, 1494 (2013).

61. Besemer, J., Lomsadze, A. & Borodovsky, M. GeneMarkS: a self-training method for prediction of gene starts in microbial genomes. Implications for finding sequence motifs in regulatory regions. Nucleic Acids Res. 29, 2607–2618 (2001).

62. Haas, B. J. et al. Improving the *Arabidopsis* genome annotation using maximal transcript alignment assemblies. Nucleic Acids Res. 31, 5654–5666 (2003).

63. Burge, C. B. & Karlin, S. Prediction of complete gene structures in human genomic DNA. J. Mol. Biol. 268, 78–94 (1997).

64. Stanke, M., Schoffmann, O., Morgenstern, B. & Waack, S. Gene prediction in eukaryotes with a generalized hidden Markov model that uses hints from external sources. BMC Bioinformatics 7, 62 (2006).

65. Korf, I. F. Gene finding in novel genomes. BMC Bioinformatics 5, 59 (2004).

66. Keilwagen, J., Hartung, F., Paulini, M., Twardziok, S. O. & Grau, J. Combining RNA-seq data and homology-based gene prediction for plants, animals and fungi. BMC Bioinformatics 19, 189 (2018).

67. Allen, J. E., Pertea, M. & Salzberg, S. L. Computational gene prediction using multiple sources of evidence. Genome Res. 14, 142–148 (2003).

68. Nawrocki, E. P., Kolbe, D. L. & Eddy, S. R. Infernal 1.0: inference of RNA alignments. Bioinformatics 25, 1335–1337 (2009).

69. Lowe, T. M. & Eddy, S. R. tRNAscan-SE: a program for improved detection of transfer RNA genes in genomic sequence. Nucleic Acids Res. 25, 955–964 (1997).

70. Birney, E., Clamp, M. & Durbin, R. GeneWise and Genomewise. Genome Res. 14, 988–995 (2004).

71. She, R., Chu, S. C., Wang, K., Pei, J. & Chen, N. GenBlastA: enabling BLAST to identify homologous gene sequences. Genome Res. 19, 143–149 (2008).

72. Roach, M. J., Schmidt, S. A. & Borneman, A. R. Purge Haplotigs: allelic contig reassignment for third-gen diploid genome assemblies. BMC Bioinformatics 19.

73. Bolger, A. M., Lohse, M. & Usadel, B. Trimmomatic: a flexible trimmer for Illumina sequence data. Bioinformatics 30, 2114–2120 (2014).

74. Li, H. Aligning sequence reads, clone sequences and assembly contigs with BWA-MEM. arXiv preprint 1303.3997 (2013).

75. Garrison, E. & Marth, G. T. Haplotype-based variant detection from short-read sequencing. 1207.3907 (2012).

76. Purcell, S. et al. PLINK: a tool set for whole-genome association and population-based linkage analyses. Am. J. Hum. Genet. 81, 559–575 (2007).

77. Zhou, X. & Stephens, M. Genome-wide efficient mixed-model analysis for association studies. Nat. Genet. 44, 821–824 (2012).

78. Quinlan, A. R. & Hall, I. M. BEDTools: a flexible suite of utilities for comparing genomic features. Bioinformatics 26, 841–842 (2010).

79. Sakai, W. S. Simple method for differential staining of paraffin embedded plant material using toluidine blue O. Stain Technology 48, 247–249 (1973).

80. Brunner, A. M., Yakovlev, I. A. & Strauss, S. H. Validating internal controls for quantitative plant gene expression studies. BMC Plant Biol. 4, 14 (2004).

81. Livak, K. J. & Schmittgen, T. D. Analysis of relative gene expression data using real-time quantitative PCR and the 2^-ΔΔCT^ method. Methods 25, 402–408 (2001).

82. Dobin, A. et al. STAR: ultrafast universal RNA-seq aligner. Bioinformatics 29, 15–21 (2013).

83. Anders, S. & Huber, W. Differential expression analysis for sequence count data. Genome Boil. 11, 1–12 (2010).

84. Yoo, S.-D., Cho, Y.-H. & Sheen, J. *Arabidopsis* mesophyll protoplasts: a versatile cell system for transient gene expression analysis. Nat. Protoc. 2, 1565 (2007).

85. Karimi, M., Inzé, D. & Depicker, A. GATEWAY™ vectors for Agrobacterium-mediated plant transformation. Trends Plant Sci. 7, 193–195 (2002).

86. Ke, S.-H. & Madison, E. L. Rapid and efficient site-directed mutagenesis by single-tube ‘megaprimer’PCR method. Nucleic Acids Res.25, 3371–3372 (1997).

87. Ebert, M. S. & Sharp, P. A. MicroRNA sponges: progress and possibilities. Curr. Biol. 20, R858–R861 (2010).

88. Clough, S. J. & Bent, A. F. Floral dip: a simplified method for *Agrobacterium*-mediated transformation of *Arabidopsis thaliana*. Plant J. 16, 735–743 (1998).

89. Smyth, D. R., Bowman, J. L. & Meyerowitz, E. M. Early flower development in *Arabidopsis*. Plant Cell 2, 755–767 (1990).

