## Supplementary material for "Two antagonistic effect genes mediate separation of sexes in a fully dioecious plant": Supplemeantal Tables

Supplementary Table 1. Primers used in this study.

| Primer | Forward primer sequence | Reverse primer sequence | Application |
|---|---|---|---|
| N025 | CCCCCTTTAAATTTTACTGTAG | TTTTCTTGTTGAGTTGACGTT | Fine mapping |
| N062 | TCCTGAATAGTACGCTACACA | ATTGATTGAAGGGCATACATA | Fine mapping |
| N084 | CTGCTACAACTCCTCCAAATA | GTGACTCAGAAAACCCATAAA | Fine mapping |
| N110 | AAAGACATTCCCTTTCTTGTC | GGGTTGTTTCCTTTTCCTT | Fine mapping |
| N126 | TTTTTCTCTCTCACAAATGGA | CATTGACTATAATGCGGGTAG | Fine mapping |
| N194 | TACCTCTCGAGTTGAAACAGA | AATGCATGAGGCAATAGTTAC | Fine mapping |
| N209 | ATTCCAACACCAACTAACTCA | GACCCTCATTGTCATGGAT | Fine mapping |
| N362 | TACATTGAATCAGCTCCAAAC | ATTCAAAGTTCAGGGATGAAA | Fine mapping |
| N283 | ATCATCTGTTTTTACGGTTCA | TTAATAGGGAGTTGGAGGAAG | Fine mapping |
| N293 | ATCCAAGTCCATTTTTCTTTC | CCTGTCTTTCTTTCGTTTGTA | Fine mapping |
| F_841 | ATAAATCCTTACGACCGCCAG | GCCAGATCTAACTCACGCAGG | Clone of SLR |
| F_815 | AACGAGTGCTTGAGTCAGAAA | AGGAGAGAGAAAGGGTGGCTA | Clone of SLR |
| F_817 | CCACCGACAAGAGACAAAGAA | TGTCTGAACAAAACGCCATTAC | Clone of SLR |
| F_821 | TGTTGATACGGATGACCACCT | CGGGATAACCCAATAAAAAGTA | Clone of SLR |
| F_651 | AGAACTATCTCCTCCCCTGCG | CCGTGTGTTGTCTGTGTGGAG | Clone of SLR |
| F_709 | ATTGCTGATCTTTTCGGTCTTT | AAATCACTTTTGTGAATTTCAGC | Clone of SLR |
| F_655 | GAGAGTCTGGTACAGTATGCTGT | AACCACAAGTCGTTCAAGTATC | Clone of SLR |
| F_710 | GTATGACAATCCCTCTCTCCCA | ATTTCTATTAGAGAGGCCCACG | Clone of SLR |
| F_657 | TGTTGTGTCGCTTATGACCAC | CTGGCGGTTATGTGAGTATTG | Clone of SLR |
| F_661 | CCACACAATACTCACATAACCGC | TTAGCAATTATCTCCGCCTCAT | Clone of SLR |
| F_712 | AAAGAATGCCCCAAGGAGAAC | TACTTGTGTTGAGAAGAGGGGG | Clone of SLR |
| F_823 | GCTGAAGCGTGACTCGATCAG | AAAGACTTCATCTCGCACACTTG | Clone of SLR |
| F_827 | AGAAAGACGCATAACAACGGTG | GAACTGTGAGCGTCGTGTGAA | Clone of SLR |
| F_830 | AGGACATCTAAACTCCCAGCT | TTCTTGTAGCCATCAATCTGC | Clone of SLR |
| F_833 | AGAGAGCCAGTTTTTTACGAAT | CAGAAAGATAAGGAGCAACAACT | Clone of SLR |
| FERR | GTGTTTTGTTGAGGAGATTAGAC | CCCTTCTTGTCTCTCTTTCTG | Clone of *FERR* |
| Mms | TTATTGTAGAAATAAGGCCTATATTCG | AAATTTATTTATAACGATCATTATCTCTCT | Clone of *Mms* |
| P801 | GTTGAGAGGTATGTGGTAGTGC | GTGTTTTCTCTACCACCACCTC | qPCR (*MET*) |
| P806 | GAGAGAACCAGACAAATGCGG | CATCCCCAATTTTCTTAACCTC | qPCR (*TCP*) |
| P977 | TTGCTATCCCATCTGGACTTT | CTTGTTGAAGTTATCAGCCACG | qPCR (*CLC*) |
| P809 | AAACGGAAAGAGAGCATTGGA | TCATAACCTGTCATTCCTGGCA | qPCR (*FERR*) |
| P982 | TCCTTGGTGTCTGAAAGTGTC | GGGAACCTTTGAAAAGTTGAA | qPCR (*MmS*) |
| P990 | AAGTAGCAATAGAGGTGGCGA | CTCTAACCCAACAGCACATCTT | qPCR (*FERR-R*) |
| PtUBQ | GTTGATTTTTGCTGGGAAGC | GATCTTGGCCTTCACGTTGT | ubiquitin |
| FERR-infusion | GACTCTAGAGGATCCATGGCCAGCTCTTCTTCTTCC | GGAAATTCGAGCTCGGTACCTTATGCATGTTCGTCTTCTCCTGCAA | *FERR*_p2301-35S plus |
| Mms-infusion | GGGGACTCTAGAGGATCCTTATTGTAGAAATAAGGCCTATATTCG | GAAATTCGAGCTCGGTACCAAATTTATTTATAACGATCATTATCTCTCT | *Mms*_p2301-35S plus |

Supplementary Table 2. Genes annotated on SLR-X and SLR-Y, showing genes whose annotation suggests that they are XY pairs of genes.

| | X haplotype | | | | | Y haplotype | | | | |
|---|---|---|---|---|---|---|---|---|---|---|
| Pairs[a] | Start | End | Dire[b] | Gene ID | Annotation | Start | End | Dire[b] | Gene ID | Annotation |
| 1 | 6516 | 10142 | - | EVM0038001 | Putative ribonuclease H protein At1g65750 | 24808 | 28035 | - | EVM0039001 | Putative ribonuclease H protein At1g65750 |
| | | | | | | 28452 | 29411 | - | EVM0039002 | None |
| | | | | | | 34621 | 42229 | - | EVM0039003 | Putative ribonuclease H protein At1g65750 |
| | 10186 | 11322 | - | EVM0038002 | None | | | | | |
| | | | | | | 43815 | 49568 | - | EVM0039004 | None |
| | | | | | | 50846 | 53313 | - | EVM0039005 | None |
| | 13949 | 20791 | - | EVM0038003 | Membrane protein of ER body-like protein | | | | | ABSENT FROM Y 1 |
| | 20888 | 41226 | + | EVM0038004 | Adenosine deaminase-like protein | | | | | ABSENT FROM Y 2 |
| | 42680 | 43414 | - | EVM0038005 | None | | | | | ABSENT FROM Y 3 |
| | 43502 | 44999 | - | EVM0038006 | Probable serine/threonine-protein kinase PBL23 | | | | | ABSENT FROM Y 4 |
| | | | | | ABSENT FROM X 1 | 61451 | 77472 | - | EVM0039006 | Two-component response regulator ARR17 |
| | | | | | ABSENT FROM X 2 | 81909 | 82697 | + | EVM0039007 | None |
| 2 | 47938 | 51624 | + | EVM0038007 | T-complex protein 1 subunit gamma(TCP) | 87483 | 94421 | + | EVM0039008 | T-complex protein 1 subunit gamma(TCP) |
| 3 | 69860 | 73840 | + | EVM0038008 | Chloride channel protein CLC-c(CLC) | 97911 | 101936 | + | EVM0039009 | Chloride channel protein CLC-c(CLC) |
| | | | | | ABSENT FROM X 3 | 102582 | 105723 | + | EVM0039010 | Transposon |
| 4 | 80892 | 90908 | + | EVM0038009 | DNA (cytosine-5)-methyltransferase 1 | 114011 | 123903 | + | EVM0039011 | DNA (cytosine-5)-methyltransferase 1 |
| | 96020 | 99449 | + | EVM0038010 | Probable disease resistance protein At1g15890 | | | | | ABSENT FROM Y 5 |
| 5 | 100609 | 103386 | + | EVM0038011 | Calcium-dependent protein kinase 4 | 129331 | 136656 | + | EVM0039012 | Calcium-dependent protein kinase 4 |

| | | | | | | | | | | |
|---|---|---|---|---|---|---|---|---|---|---|
| | 105755 | 106240 | - | EVM0038012 | None | | | | | |
| | 109496 | 110467 | + | EVM0038013 | None | | | | | |
| | 113188 | 113559 | + | EVM0038014 | None | | | | | |
| | | | | | | 138996 | 139496 | - | EVM0039013 | None |
| | | | | | | 146409 | 146780 | + | EVM0039014 | None |
| 6 | 113610 | 113873 | + | EVM0038015 | E3 ubiquitin-protein ligase makorin | 146831 | 147094 | + | EVM0039015 | E3 ubiquitin-protein ligase makorin |
| | 115866 | 116422 | - | EVM0038016 | None | | | | | |
| | 116471 | 118068 | - | EVM0038017 | None | | | | | |
| 7 | 118349 | 126441 | + | EVM0038018 | Cleft lip and palate transmembrane protein 1 homolog | 147865 | 155841 | + | EVM0039016 | Cleft lip and palate transmembrane protein 1 homolog |
| 8 | 128479 | 133608 | - | EVM0038019 | Protein SUPPRESSOR OF GENE SILENCING 3 | 157892 | 163032 | - | EVM0039017 | Protein SUPPRESSOR OF GENE SILENCING 3 |
| 9 | 136821 | 139171 | - | EVM0038020 | None | 167821 | 168045 | - | EVM0039018 | None |
| 10 | 142264 | 143104 | + | EVM0038021 | PHD finger protein At1g33420 | 172166 | 177190 | + | EVM0039019 | PHD finger protein At1g33420 |
| | 143296 | 144731 | + | EVM0038022 | None | | | | | |
| | 145597 | 145828 | - | EVM0038023 | None | | | | | |
| | 146378 | 146602 | - | EVM0038024 | None | | | | | |
| | 150695 | 155648 | + | EVM0038025 | PHD finger protein At1g33420 | | | | | |
| 11 | 156184 | 159201 | + | EVM0038026 | Peroxidase 47 | 177652 | 180652 | + | EVM0039020 | Peroxidase 47 |
| 12 | 161059 | 164210 | - | EVM0038027 | 4,5-DOPA dioxygenase extradiol | 182505 | 185706 | - | EVM0039021 | 4,5-DOPA dioxygenase extradiol |
| 13 | 166480 | 180086 | - | EVM0038028 | Probable disease resistance protein At4g27220 | 188011 | 190086 | - | EVM0039022 | Probable disease resistance protein At4g27220 |
| | 185375 | 185461 | + | EVM0038029 | None | | | | | |
| | 185494 | 186861 | - | EVM0038030 | None | | | | | |

| | | | | | | | | | | |
|---|---|---|---|---|---|---|---|---|---|---|
| | 188133 | 190558 | - | EVM0038031 | Probable LRR receptor-like serine/threonine-protein kinase At1g07650 | | | | | ABSENT FROM Y 6 |
| | 190717 | 194232 | - | EVM0038032 | Putative disease resistance protein RGA4 | | | | | ABSENT FROM Y 7 |
| | 198390 | 201014 | + | EVM0038033 | None | | | | | |
| | 201032 | 201112 | - | EVM0038034 | None | | | | | |
| | 201595 | 204301 | - | EVM0038035 | None | | | | | |
| | 205006 | 206255 | - | EVM0038036 | None | | | | | |
| 14 | 209242 | 212680 | - | EVM0038037 | Probable disease resistance protein At4g27220 | 197721 | 202988 | - | EVM0039023 | Probable disease resistance protein At4g27220 |
| | 218494 | 219026 | - | EVM0038038 | Probable LRR receptor-like serine/threonine-protein kinase At1g07650 | | | | | ABSENT FROM Y 8 |
| 15 | 225393 | 229630 | - | EVM0038039 | Probable disease resistance protein At4g27220 | 214009 | 218246 | - | EVM0039024 | Probable disease resistance protein At4g27220 |
| | 230086 | 231963 | - | EVM0038040 | None | | | | | |
| | | | | | | 220236 | 227934 | - | EVM0039025 | None |
| 16 | 241933 | 245026 | + | EVM0038041 | Probable disease resistance protein At5g43730 | 238511 | 241603 | + | EVM0039026 | Probable disease resistance protein At5g43730 |

[a] XY gene pairs
[b] Direction

Supplementary Table 3. Genome resequencing statistics of the female and male *P. deltoides*.

| Sample ID | Sex | Read Base | Read Number | Coverage |
|---|---|---|---|---|
| 36-1 | Female | 11,649,192,900 | 38,830,643 | 27.09 |
| 40-1 | Female | 11,168,084,700 | 37,226,949 | 25.97 |
| 42-4 | Female | 12,287,930,700 | 40,959,769 | 28.58 |
| 47-4 | Female | 12,636,568,200 | 42,121,894 | 29.39 |
| 48-5 | Female | 11,102,390,400 | 37,007,968 | 25.82 |
| 50-5 | Female | 14,602,650,900 | 48,675,503 | 33.96 |
| 52-1 | Female | 11,245,898,400 | 37,486,328 | 26.15 |
| 60-8 | Female | 13,177,485,900 | 43,924,953 | 30.65 |
| 63-1 | Female | 11,296,434,900 | 37,654,783 | 26.27 |
| 66-5 | Female | 11,477,944,500 | 38,259,815 | 26.69 |
| 70-5 | Female | 10,964,830,500 | 36,549,435 | 25.50 |
| 72-6 | Female | 11,179,380,300 | 37,264,601 | 26.00 |
| 74-4 | Female | 11,733,299,700 | 39,110,999 | 27.29 |
| 77-1 | Female | 10,556,599,800 | 35,188,666 | 24.55 |
| 78-4 | Female | 11,534,989,800 | 38,449,966 | 26.83 |
| 80-4 | Female | 11,815,474,500 | 39,384,915 | 27.48 |
| 83-3 | Female | 14,071,658,400 | 46,905,528 | 32.72 |
| 87-6 | Female | 10,132,395,000 | 33,774,650 | 23.56 |
| 88-2 | Female | 11,882,076,600 | 39,606,922 | 27.63 |
| 90-5 | Female | 13,220,877,300 | 44,069,591 | 30.75 |
| 92-7 | Female | 9,888,900,000 | 32,963,000 | 23.00 |
| 93-1 | Female | 11,695,936,500 | 38,986,455 | 27.20 |
| 96-7 | Female | 11,972,586,300 | 39,908,621 | 27.84 |
| 97-3 | Female | 11,297,386,500 | 37,657,955 | 26.27 |
| 98-5 | Female | 15,165,401,700 | 50,551,339 | 35.27 |
| 22-1 | Female | 12,758,267,700 | 42,527,559 | 29.67 |
| 24-1 | Female | 11,006,373,300 | 36,687,911 | 25.60 |
| 31-1 | Female | 11,898,806,700 | 39,662,689 | 27.67 |
| 28-3 | Female | 11,370,264,300 | 37,900,881 | 26.44 |
| 7-3 | Female | 11,660,032,200 | 38,866,774 | 27.12 |
| 11-1 | Female | 11,459,820,900 | 38,199,403 | 26.65 |
| 100-2 | Female | 12,408,186,000 | 41,360,620 | 28.86 |
| 101-5 | Female | 11,580,759,900 | 38,602,533 | 26.93 |
| 102-3 | Female | 11,684,571,900 | 38,948,573 | 27.17 |
| 103-3 | Female | 11,444,420,700 | 38,148,069 | 26.61 |
| 104-1 | Female | 11,093,350,500 | 36,977,835 | 25.80 |
| 61-18 | Female | 12,642,730,800 | 42,142,436 | 29.40 |
| S3230 | Female | 14,264,241,300 | 47,547,471 | 33.17 |
| 81-22 | Female | 11,090,319,000 | 36,967,730 | 25.79 |
| 86-13 | Female | 11,822,777,400 | 39,409,258 | 27.49 |

| | | | | |
|---|---|---|---|---|
| 89-18 | Female | 11,622,264,300 | 38,740,881 | 27.03 |
| 94-27 | Female | 11,794,462,200 | 39,314,874 | 27.43 |
| 95-30 | Female | 11,465,142,300 | 38,217,141 | 26.66 |
| S3016 | Female | 12,946,626,300 | 43,155,421 | 30.11 |
| S3109 | Female | 12,148,255,800 | 40,494,186 | 28.25 |
| S3229 | Female | 13,144,587,900 | 43,815,293 | 30.57 |
| S3107 | Female | 12,787,382,100 | 42,624,607 | 29.74 |
| S3700 | Female | 15,085,499,100 | 50,284,997 | 35.08 |
| S3406 | Female | 11,765,872,500 | 39,219,575 | 27.36 |
| 31-4 | Male | 12,545,672,100 | 41,818,907 | 29.18 |
| 35-2 | Male | 11,043,030,000 | 36,810,100 | 25.68 |
| 36-3 | Male | 11,442,942,900 | 38,143,143 | 26.61 |
| 45-1 | Male | 13,935,390,600 | 46,451,302 | 32.41 |
| 47-2 | Male | 11,552,744,700 | 38,509,149 | 26.87 |
| 48-1 | Male | 11,386,119,000 | 37,953,730 | 26.48 |
| 51-1 | Male | 12,487,357,200 | 41,624,524 | 29.04 |
| 60-2 | Male | 11,990,539,200 | 39,968,464 | 27.88 |
| 62-3 | Male | 11,821,741,500 | 39,405,805 | 27.49 |
| 63-5 | Male | 11,965,007,100 | 39,883,357 | 27.83 |
| 70-1 | Male | 11,459,258,100 | 38,197,527 | 26.65 |
| 72-2 | Male | 12,000,861,900 | 40,002,873 | 27.91 |
| 74-1 | Male | 14,045,237,100 | 46,817,457 | 32.66 |
| 77-10 | Male | 11,668,235,700 | 38,894,119 | 27.14 |
| 78-2 | Male | 11,339,813,400 | 37,799,378 | 26.37 |
| 86-6 | Male | 11,183,043,900 | 37,276,813 | 26.01 |
| 87-5 | Male | 11,769,492,300 | 39,231,641 | 27.37 |
| 88-7 | Male | 11,964,074,100 | 39,880,247 | 27.82 |
| 93-4 | Male | 11,028,985,200 | 36,763,284 | 25.65 |
| 95-12 | Male | 11,846,649,600 | 39,488,832 | 27.55 |
| 96-10 | Male | 13,193,655,900 | 43,978,853 | 30.68 |
| 28-2 | Male | 12,408,126,600 | 41,360,422 | 28.86 |
| 24-3 | Male | 11,172,411,300 | 37,241,371 | 25.98 |
| 22-4 | Male | 11,966,299,200 | 39,887,664 | 27.83 |
| 100-4 | Male | 11,177,231,400 | 37,257,438 | 25.99 |
| 101-3 | Male | 10,894,545,900 | 36,315,153 | 25.34 |
| 102-9 | Male | 11,939,396,100 | 39,797,987 | 27.77 |
| 103-9 | Male | 11,884,926,600 | 39,616,422 | 27.64 |
| 104-2 | Male | 12,839,783,400 | 42,799,278 | 29.86 |
| 131-3 | Male | 13,945,134,900 | 46,483,783 | 32.43 |
| S3301 | Male | 11,829,934,500 | 39,433,115 | 27.51 |
| 65-18 | Male | 11,610,252,900 | 38,700,843 | 27.00 |
| 66-16 | Male | 11,715,514,500 | 39,051,715 | 27.25 |
| 81-30 | Male | 11,411,807,700 | 38,039,359 | 26.54 |

| | | | | |
|---|---|---|---|---|
| 89-28 | Male | 11,875,062,600 | 39,583,542 | 27.62 |
| 94-28 | Male | 11,835,697,200 | 39,452,324 | 27.52 |
| S3101 | Male | 13,373,461,500 | 44,578,205 | 31.10 |
| S3201 | Male | 14,262,937,800 | 47,543,126 | 33.17 |
| S3244 | Male | 11,973,510,900 | 39,911,703 | 27.85 |
| S3261 | Male | 13,581,076,200 | 45,270,254 | 31.58 |
| S3415 | Male | 11,210,397,600 | 37,367,992 | 26.07 |
| S3702 | Male | 14,024,592,000 | 46,748,640 | 32.62 |
| S3804 | Male | 13,570,024,800 | 45,233,416 | 31.56 |
| S3236 | Male | 11,486,967,600 | 38,289,892 | 26.71 |
| S3239 | Male | 14,794,821,300 | 49,316,071 | 34.41 |
| S3240 | Male | 13,313,584,500 | 44,378,615 | 30.96 |

Supplementary Table 4. Summary of genome regions containing SNPs with fully sex-linked genotype configurations.

| Chromosome | Start of region | End of region | Number of SNPs. | Number of genes |
|---|---|---|---|---|
| XIX SLR of XY pair | 120,289 | 161,988 | 315 | 3 |
| XIX PAR of XY pair | 17,899,199 | 17,909,028 | 78 | 1 (FERR) |
| IX (autosome) | 7,729,222 | 7,730,262 | 27 | 1 (HEMA1) |
| I | 50,047,869 | 50,053,274 | 5 | 2 |
| XVIII | 4,573,272 | 4,573,288 | 2 | 1 |
| I | 22,376,091 | — | 1 | 0 |
| Contig01665 | 13,437 | 16,183 | 7 | 2 |
| Total | 435 | 10 | | |

Supplementary Table 5. Sequence homology analysis for sequence containing SEMSs.

| Location of SEMSs | Length of query sequence | Hit start in the YHF | Hit end in the YHF | Identity (%) | E Value |
|---|---|---|---|---|---|
| PAR | 10,030 | 61,590 | 61,418 | 92.49 | 6.42E-63 |
| PAR | 10,030 | 61,683 | 61,588 | 93.75 | 8.67E-32 |
| PAR | 10,030 | 61,767 | 63,351 | 87.59 | 0 |
| PAR | 10,030 | 66,854 | 63,357 | 88.95 | 0 |
| PAR | 10,030 | 66,965 | 67,466 | 88.76 | 7.11E-172 |
| PAR | 10,030 | 67,601 | 68,223 | 90.21 | 0 |
| PAR | 10,030 | 68,416 | 69,127 | 92.61 | 0 |
| PAR | 10,030 | 68,460 | 66,820 | 85.42 | 0 |
| PAR | 10,030 | 69,398 | 69,226 | 93.06 | 1.38E-64 |
| PAR | 10,030 | 69,425 | 73,224 | 88.61 | 0 |
| PAR | 10,030 | 74,662 | 74,993 | 94.29 | 2.67E-141 |
| PAR | 10,030 | 78,458 | 75,021 | 89.37 | 0 |
| PAR | 10,030 | 78,712 | 79,523 | 92.16 | 0 |
| PAR | 10,030 | 79,516 | 80,026 | 92.25 | 0 |
| Chr_I | 5,606 | 49,675 | 47,836 | 89.24 | 0 |
| Chr_I | 5,606 | 49,856 | 49,671 | 89.25 | 1.00E-58 |
| Chr_I | 201 | 53,477 | 53,286 | 89.45 | 1.45E-63 |
| Chr_IX | 1,241 | 73,287 | 74,402 | 89.26 | 0 |
| Chr_XVIII | 217 | 104,531 | 104,747 | 91.71 | 9.29E-81 |
| Contig01665 | 2,947 | 106,608 | 103,675 | 93.49 | 0 |

Supplementary Table 6. miRNA binding sites in *MmS* transcripts, and the miRNAs they target.

| Pair | microRNA | | Alignment | |
|---|---|---|---|---|
| | *MmS* | 67 | AAUGAAUGCUG-AUUUAAUGA | 86 |
| 1 |  |  | :: ::.:::: :.:.::::: |  |
| | miR397a | 21 | GUAGUUGCGACGUGAGUUACU | 1 |
| | *MmS* | 237 | GGGACCCGACUUCUCGCGUGC | 257 |
| 2 |  |  | : :::::.:.: :.::: |  |
| | miR160e | 21 | GGACGAGCUGAGGGGAGUACG | 1 |
| | *MmS* | 285 | UUGCCCCCUUGCUUCU-UCA | 303 |
| 3 |  |  | ::: ::::. ::::: ::: |  |
| | miR156k | 20 | CACGAGGGAGAGAAGACAGU | 1 |
| | *MmS* | 293 | UUGCUUCUUCAAAUUGUUUAA | 313 |
| 4 |  |  | : :.::.::::.. ::.: |  |
| | miR395a | 21 | CUCAAGGAGGUUUGGGAAGUC | 1 |
| | *MmS* | 469 | GAAUUCUUUGUCUCUUGUCGG | 489 |
| 5 |  |  | : .:::.:.::: .::::.. |  |
| | miR156g | 21 | CACGAGAGAUAGAAGACAGUU | 1 |
| | *MmS* | 750 | GUUCUAACGUCUAUGUAAAAA | 770 |
| 6 | | | : .:: : .::.:.:.: ::: | |
| | miR403c | 21 | CCGGAGUCUAGGUGCGUGUUU | 1 |
| | *MmS* | 1538 | GGCUGCUCUUCCUCCUCUCUG | 1558 |
| 7 |  |  | :::::: ::: ::.:.: |  |
| | miR408 | 21 | GGUACGAGACGGACAAGGGGC | 1 |
| | *MmS* | 1701 | UGCAGAGGCGUUGCUUCUUCU | 1721 |
| 8 |  |  | :::: :: .:::::::.: |  |
| | miR164a | 21 | ACGUGCACGGGACGAAGAGGU | 1 |
| | *MmS* | 1768 | UAGCAGCUUCAUGGAGGUGCA | 1788 |
| 9 |  |  | ::::: :::: .::.: : |  |
| | miR172a | 21 | UACGUCGUAGUAGUUCUAAGA | 1 |
| | *MmS* | 2046 | GGUGUGUUUGUAUUUCCCUCC | 2066 |
| 10 |  |  | .:::: :: .: :: :::: |  |
| | miR398c | 21 | GUACACUAAAGUCCAGCGAGG | 1 |
| | *MmS* | 2082 | CCUUUCCUUCUCUUCUCUCC | 2101 |
| 11 |  |  | .:::.:::::::: :: |  |
| | miR156k | 20 | CACGAGGGAGAGAAGACAGU | 1 |
| | *MmS* | 2098 | CUCCGUUCACUCUUCUCUCGUU | 2119 |
| 12 |  |  | :.::: : :.::::.:.: |  |
| | miR482c | 22 | UCAGUAAGGGCGGAGAGGGUAU | 1 |
| | *MmS* | 2157 | UCGUUCCUUCUUUUGUGUUU | 2176 |
| 13 |  |  | :.:::.:::.:: :::. |  |
| | miR156k | 20 | CACGAGGGAGAGAAGACAGU | 1 |

| | | | | |
|---|---|---|---|---|
| 14 | *MmS* | 2214 | UACCCUCUCACAGUAGCUUUCU | 2235 |
|  |  |  | . :.:.::::. :: ::::. |  |
| | miR477c | 22 | GUUUGGGGGUGUUUUCCAAAGG | 1 |
| 15 | *MmS* | 2225 | AGUAGCUUUCUUUGUUUCAAA | 2245 |
|  |  |  | . ::::..:::.. :.:::: |  |
| | miR159a | 21 | AUCUCGAGGGAAGUUAGGUUU | 1 |
| 16 | *MmS* | 2225 | AGUAGCUUUCUUUGUUUCAA | 2244 |
|  |  |  | : ::::..:::.. :.::: |  |
| | miR319a | 20 | CCCUCGAGGGAAGUCAGGUU | 1 |
| 17 | *MmS* | 2507 | GGCUUUAGGCAGUGUGUGGAG | 2527 |
|  |  |  | ::.:.: :. ::::::. |  |
| | miR396a | 21 | GUCAAGUUCUUUCGACACCUU | 1 |
| 18 | *MmS* | 2515 | GCAGUGUGUGGAG-AGAGAGAG | 2535 |
| | | | .:: :::::: :: ::.:. | |
| | miR472a | 22 | CCCUACCCACCUCAUCCCUUUU | 1 |
| 19 | *MmS* | 2526 | AGAGAGAGAGAGAUGAACAG | 2545 |
| | | | : :.:::::::. ::::: | |
| | miR394a | 20 | AAUGUUUCUCUCUGGUUGUC | 1 |
| 20 | *MmS* | 2627 | CGGGCAAUACAAAUAUUGUUG | 2647 |
|  |  |  | :.:::::: :: .:: :.:. |  |
| | miR169y | 21 | GUCCGUUAAGUAGGUACCGAU | 1 |
| 21 | *MmS* | 2657 | UACCAACAAGUAGGAGAAAUAUUA | 2680 |
| | | | : :: : ::.::::::: :. ::: | |
| | miR478h | 24 | AGGGAUUUUUAUCCUCUGUGCAAU | 1 |
| 22 | *MmS* | 2819 | GAUAAUAAAUGUGUCGUGUAA | 2839 |
|  |  |  | :::: :::: :. ::::: |  |
| | miR475a | 21 | GAAUUAGUUACCCGUGACAUU | 1 |

Supplementary Table 7. List of samples used for sequencing in this study.

| Sample | Tissue | Library_ID | Application |
|---|---|---|---|
| *P. deltoides* ♀ | Flower bud | T2 (June 18)_bud_1_F | lncRNA-seq |
| *P. deltoides* ♀ | Flower bud | T2 (June 18)_bud_2_F | lncRNA-seq |
| *P. deltoides* ♀ | Flower bud | T2 (June 18)_bud_3_F | lncRNA-seq |
| *P. deltoides* ♀ | Flower bud | T3 (July 3)_bud_1_F | lncRNA-seq |
| *P. deltoides* ♀ | Flower bud | T3 (July 3)_bud_2_F | lncRNA-seq |
| *P. deltoides* ♀ | Flower bud | T3 (July 3)_bud_3_F | lncRNA-seq |
| *P. deltoides* ♀ | Flower bud | T4 (July 18)_bud_1_F | lncRNA-seq |
| *P. deltoides* ♀ | Flower bud | T4 (July 18)_bud_2_F | lncRNA-seq |
| *P. deltoides* ♀ | Flower bud | T4 (July 18)_bud_3_F | lncRNA-seq |
| *P. deltoides* ♀ | Descaled flower bud | T5 (August 3)_bud_1_F | lncRNA-seq |
| *P. deltoides* ♀ | Descaled flower bud | T5 (August 3)_bud_2_F | lncRNA-seq |
| *P. deltoides* ♀ | Descaled flower bud | T5 (August 3)_bud_3_F | lncRNA-seq |
| *P. deltoides* ♀ | Descaled flower bud | T8 (December 1)_bud_1_F | lncRNA-seq |
| *P. deltoides* ♀ | Descaled flower bud | T8 (December 1)_bud_2_F | lncRNA-seq |
| *P. deltoides* ♀ | Descaled flower bud | T8 (December 1)_bud_3_F | lncRNA-seq |
| *P. deltoides* ♂ | Flower bud | T2 (June 18)_bud_1_M | lncRNA-seq |
| *P. deltoides* ♂ | Flower bud | T2 (June 18)_bud_2_M | lncRNA-seq |
| *P. deltoides* ♂ | Flower bud | T2 (June 18)_bud_3_M | lncRNA-seq |
| *P. deltoides* ♂ | Flower bud | T3 (July 3)_bud_1_M | lncRNA-seq |
| *P. deltoides* ♂ | Flower bud | T3 (July 3)_bud_2_M | lncRNA-seq |
| *P. deltoides* ♂ | Flower bud | T3 (July 3)_bud_3_M | lncRNA-seq |
| *P. deltoides* ♂ | Flower bud | T4 (July 18)_bud_1_M | lncRNA-seq |
| *P. deltoides* ♂ | Flower bud | T4 (July 18)_bud_2_M | lncRNA-seq |
| *P. deltoides* ♂ | Flower bud | T4 (July 18)_bud_3_M | lncRNA-seq |
| *P. deltoides* ♂ | Descaled flower bud | T5 (August 3)_bud_1_M | lncRNA-seq |
| *P. deltoides* ♂ | Descaled flower bud | T5 (August 3)_bud_2_M | lncRNA-seq |
| *P. deltoides* ♂ | Descaled flower bud | T5 (August 3)_bud_3_M | lncRNA-seq |
| *P. deltoides* ♂ | Descaled flower bud | T8 (December 1)_bud_1_M | lncRNA-seq |
| *P. deltoides* ♂ | Descaled flower bud | T8 (December 1)_bud_2_M | lncRNA-seq |
| *P. deltoides* ♂ | Descaled flower bud | T8 (December 1)_bud_3_M | lncRNA-seq |
| *P. deltoides* ♀ | Flower bud | T2 (June 18)_bud_1_F | small RNA |
| *P. deltoides* ♀ | Flower bud | T2 (June 18)_bud_2_F | small RNA |
| *P. deltoides* ♀ | Flower bud | T2 (June 18)_bud_3_F | small RNA |
| *P. deltoides* ♀ | Descaled flower bud | T8 (December 1)_bud_1_F | small RNA |
| *P. deltoides* ♀ | Descaled flower bud | T8 (December 1)_bud_2_F | small RNA |
| *P. deltoides* ♀ | Descaled flower bud | T8 (December 1)_bud_3_F | small RNA |
| *P. deltoides* ♀ | Descaled flower bud | T9 (January 15)_bud_1_F | small RNA |
| *P. deltoides* ♀ | Descaled flower bud | T9 (January 15)_bud_2_F | small RNA |
| *P. deltoides* ♀ | Descaled flower bud | T9 (January 15)_bud_3_F | small RNA |
| *P. deltoides* ♂ | Flower bud | T2 (June 18)_bud_1_M | small RNA |

| | | | |
|---|---|---|---|
| *P. deltoides* ♂ | Flower bud | T2 (June 18)_bud_2_M | small RNA |
| *P. deltoides* ♂ | Flower bud | T2 (June 18)_bud_3_M | small RNA |
| *P. deltoides* ♂ | Descaled flower bud | T8 (December 1)_bud_1_M | small RNA |
| *P. deltoides* ♂ | Descaled flower bud | T8 (December 1)_bud_2_M | small RNA |
| *P. deltoides* ♂ | Descaled flower bud | T8 (December 1)_bud_3_M | small RNA |
| *P. deltoides* ♂ | Descaled flower bud | T9 (January 15)_bud_1_M | small RNA |
| *P. deltoides* ♂ | Descaled flower bud | T9 (January 15)_bud_2_M | small RNA |
| *P. deltoides* ♂ | Descaled flower bud | T9 (January 15)_bud_3_M | small RNA |
| *P. deltoides* ♀ | Flower bud | T3 (July 3)_bud_1_F | DNA methylation |
| *P. deltoides* ♀ | Flower bud | T3 (July 3)_bud_2_F | DNA methylation |
| *P. deltoides* ♀ | Flower bud | T3 (July 3)_bud_3_F | DNA methylation |
| *P. deltoides* ♀ | Descaled flower bud | T8 (December 1)_bud_1_F | DNA methylation |
| *P. deltoides* ♀ | Descaled flower bud | T8 (December 1)_bud_2_F | DNA methylation |
| *P. deltoides* ♀ | Descaled flower bud | T8 (December 1)_bud_3_F | DNA methylation |
| *P. deltoides* ♀ | Descaled flower bud | T9 (January 15)_bud_1_F | DNA methylation |
| *P. deltoides* ♀ | Descaled flower bud | T9 (January 15)_bud_2_F | DNA methylation |
| *P. deltoides* ♀ | Descaled flower bud | T9 (January 15)_bud_3_F | DNA methylation |
| *P. deltoides* ♂ | Flower bud | T3 (July 3)_bud_1_M | DNA methylation |
| *P. deltoides* ♂ | Flower bud | T3 (July 3)_bud_2_M | DNA methylation |
| *P. deltoides* ♂ | Flower bud | T3 (July 3)_bud_3_M | DNA methylation |
| *P. deltoides* ♂ | Descaled flower bud | T8 (December 1)_bud_1_M | DNA methylation |
| *P. deltoides* ♂ | Descaled flower bud | T8 (December 1)_bud_2_M | DNA methylation |
| *P. deltoides* ♂ | Descaled flower bud | T8 (December 1)_bud_3_M | DNA methylation |
| *P. deltoides* ♂ | Descaled flower bud | T9 (January 15)_bud_1_M | DNA methylation |
| *P. deltoides* ♂ | Descaled flower bud | T9 (January 15)_bud_2_M | DNA methylation |
| *P. deltoides* ♂ | Descaled flower bud | T9 (January 15)_bud_3_M | DNA methylation |

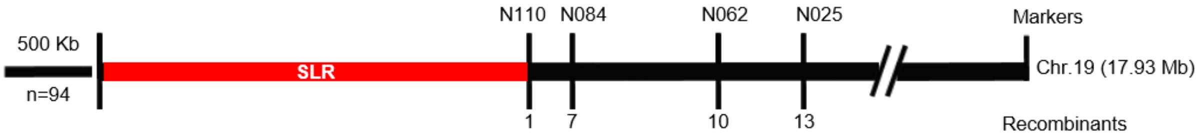


**Supplementary Figure 1. Initial mapping of sex-linked locus.** A small population of 94 *P. deltoides* individuals was screened using SSR markers.

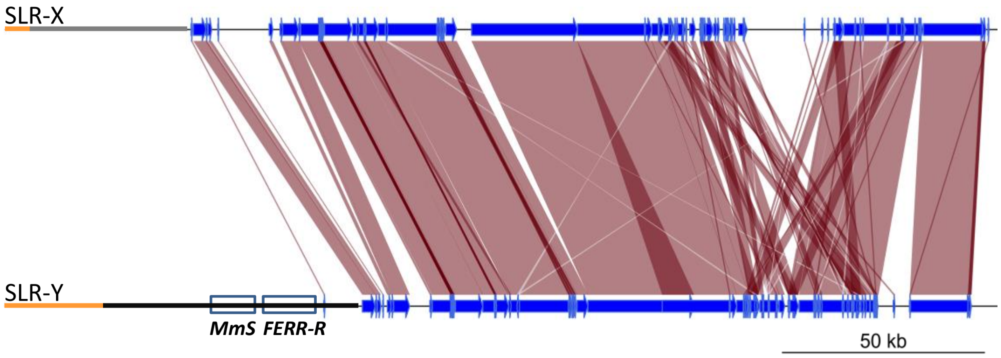


**Supplementary Figure 2. Alignment of sex-linked regions of two haplotypes.**

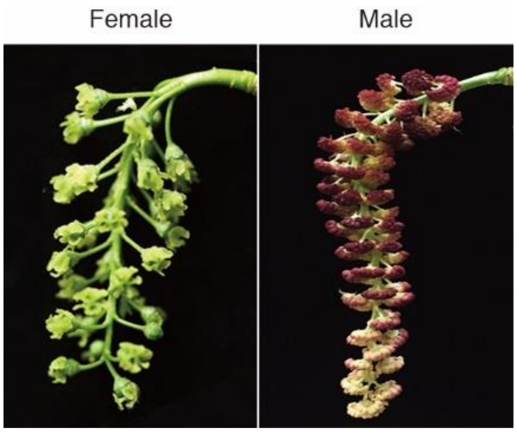


**Supplementary Figure 3. Female and male flowers of *P. deltoides*.**

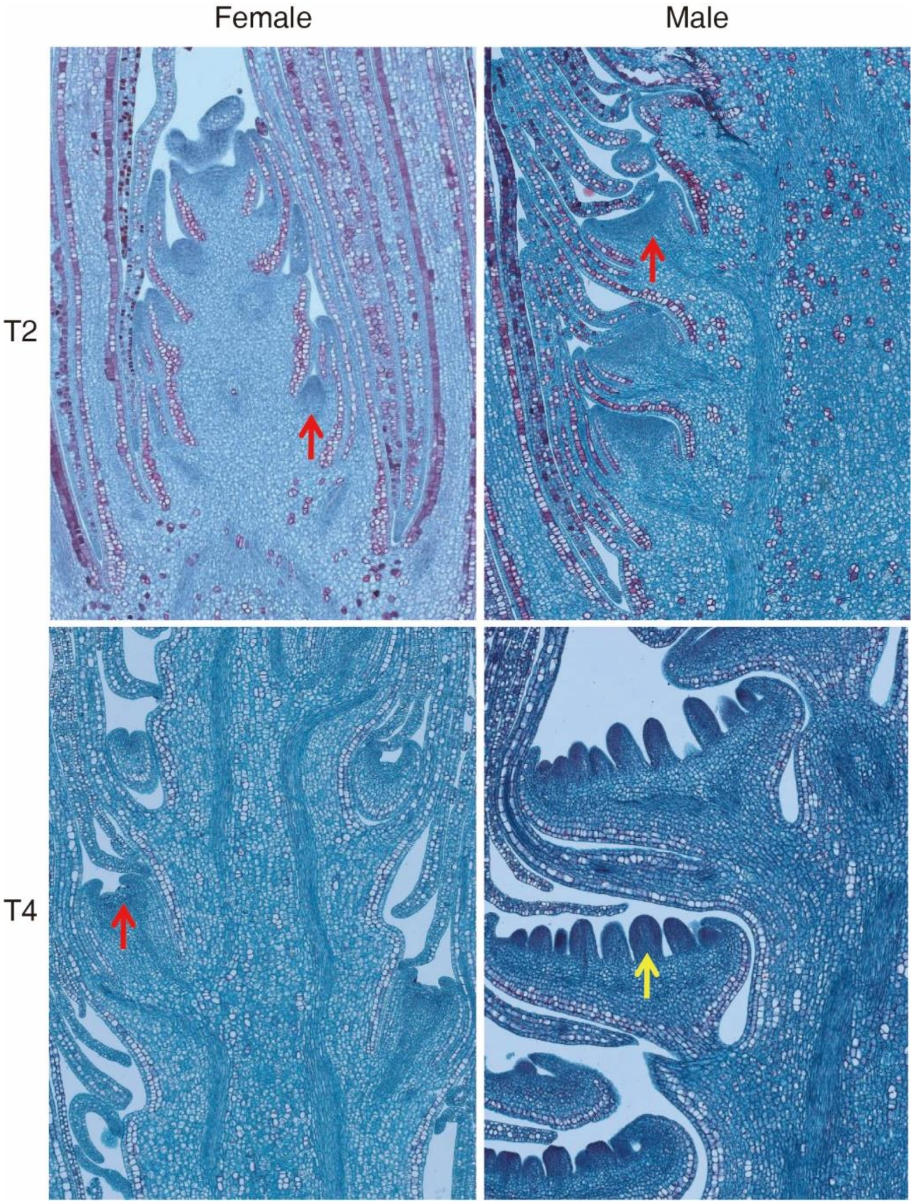


**Supplementary Figure 4. The longitudinal section of the male and female inflorescences at T2 and T4.** T2 and T4 represent June 18 and July 18 respectively. The red arrows point to the florets primordium, whereas the yellow arrow denotes the anther primordium.

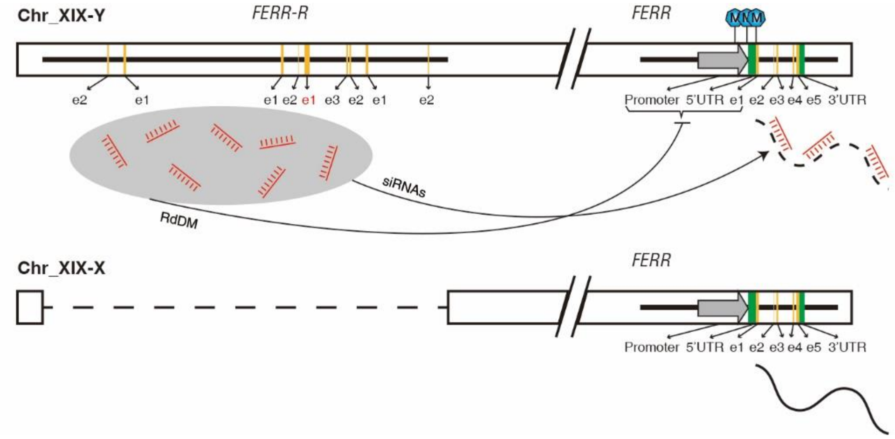


**Supplementary Figure 5. The proposed model for *FERR-R*.** e1-e5 represent exons of the *FERR* gene. RdDM indicates RNA-directed DNA methylation, and "siRNAs" indicates siRNA-guided mRNA cleavage. XX females do not undergo the cleavage, and female structures develop. XY males express the Y-linked *FERR-R* gene, and undergo the cleavage, suppressing development of female flowers.

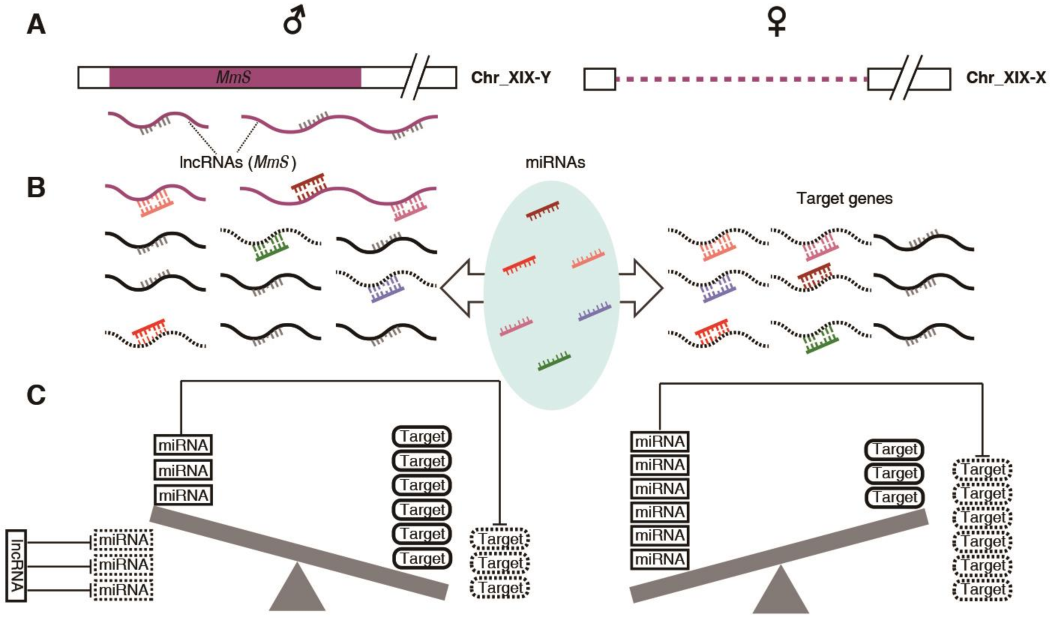


**Supplementary Figure 6. The proposed working hypothesis for the action of *MmS*.** (**A**) The Y-specific *MmS* transcribes lncRNAs, which remove miRNAs specifically in males (**B**). (**C**) Target genes are therefore expressed more highly in individuals that carry this gene, which develop male flowers (left part of the figure) than in females (on the right in the figure).

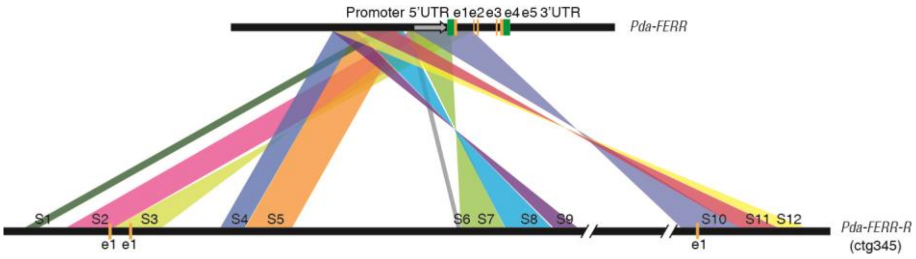


**Supplementary Figure 7. Origination of *FERR-R* in *P. davidiana*.** e1-e5 represent exons of *FERR* in *P. davidiana*. S1-S12 represent the duplicated segments between *FERR-R* and *FERR* in *P. davidiana.*

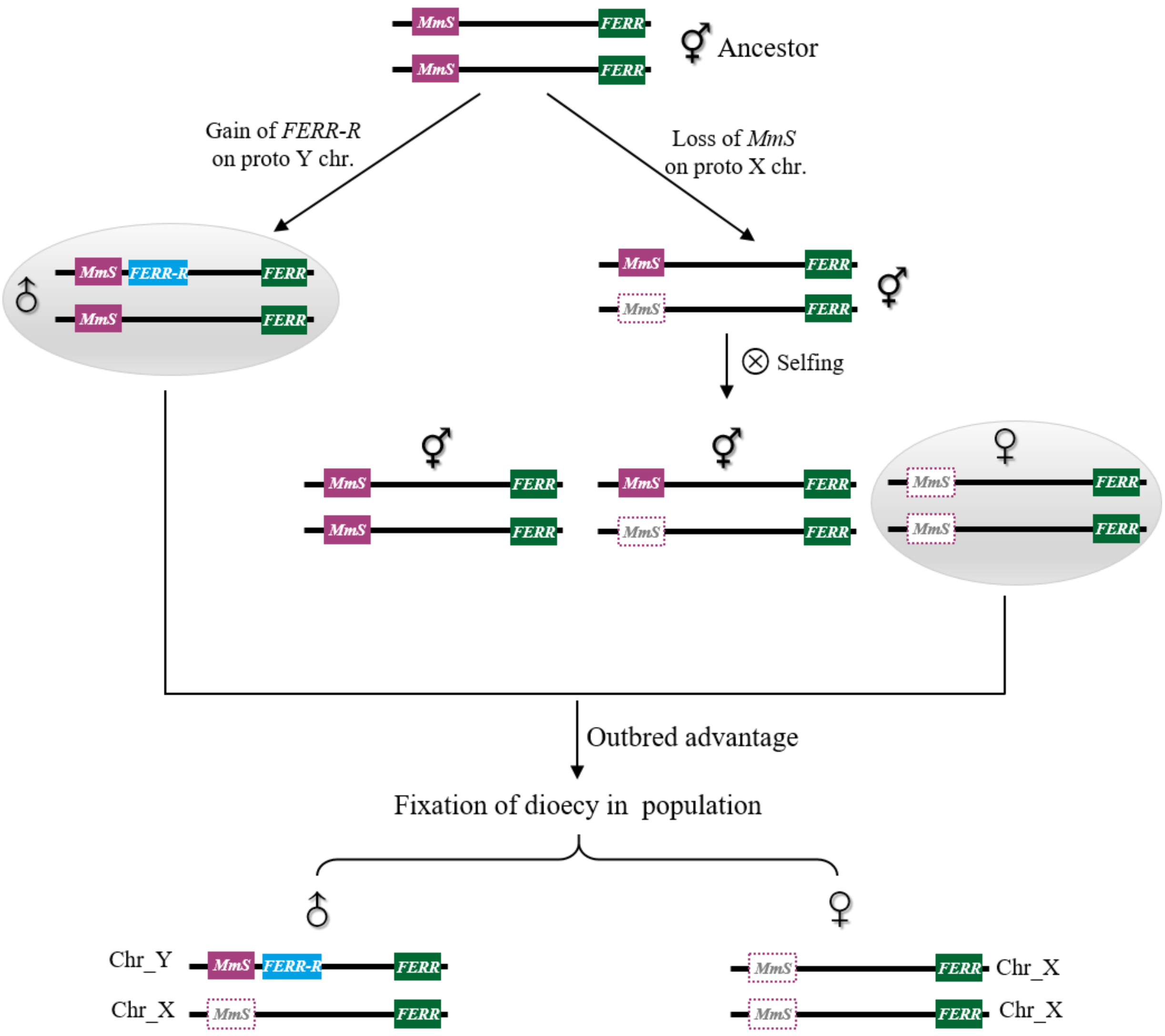


**Supplementary Figure 8. Diagram of a two-gene model for the evolution of dioecy in *P. deltoids*.** In this model, the putative hermaphorditic ancestor bears the *MmS* and *FERR* orthologs. Females could have arisen by loss of the *MmS* gene on a proto X-chromosome, which would produce a YHF region in an SLR-Y. In the absence of *FERR-R*, this genotype cannot produce male flowers. Subsequently, males could have arisen by a duplication of *FERR* that produced the *FERR-R* sequence that acts as a female-suppressor mutation. Driving by the outbred advantage, dioecy will be fixed in the population. This duplication must have occurred very physically close to the *MmS* gene to ensure the male descendants inherit *FERR-R* and *MmS* as a single unit, thereby to establish the XY system in the extant *P. deltiodes* population.
